# Hepatic conversion of acetyl-CoA to acetate plays crucial roles in energy stresses

**DOI:** 10.1101/2023.03.23.533947

**Authors:** Jinyang Wang, Yaxin Wen, Wentao Zhao, Yan Zhang, Furong Lin, Cong Ouyang, HuiHui Wang, Lizheng Yao, Huanhuan Ma, Yue Zhuo, Huiying Huang, Xiulin Shi, Liubin Feng, Donghai Lin, Bin Jiang, Qinxi Li

## Abstract

Accumulating evidences indicate that acetate is increased in energy stresses such as diabetes mellitus and prolonged starvation. However, it is largely unknown how and where acetate is produced and what is its biological significance. We observed overproduction of acetate in an amount comparable to ketone bodies in patients and mice with diabetes or starvation. Mechanistically, ACOT 12&8 are dramatically upregulated in liver to convert FFA-derived acetyl-CoA to acetate and CoA. This conversion not only provides large amount of acetate which fuels brain preferentially rather than muscle, but also recycles CoA which is required for sustained fatty acid oxidation and ketogenesis. Taken together, we suggest that acetate is an emerging novel “ketone body” and may be used as a parameter to evaluate the progression of energy stress in the future.

## Introduction

Homeostasis disorder of energy metabolism associated with emergency status such as untreated diabetes mellitus, prolonged starvation and ischemic heart/brain diseases leads to serious threats to human health (Field et al., 2001; Galgani and Ravussin, 2008; Martinic and von Herrath, 2008; Must et al., 1999). In response to such disorder, the metabolic patterns of multiple organs have to be remodelled to rescue the imbalance and bring whole organism through the crises (Denechaud et al., 2008; Fruhbeck et al., 2001; Goldberg et al., 2018; Hirai et al., 2021; Meier and Gressner, 2004; Nishimoto et al., 2016; Palikaras et al., 2015; Russell and Cook, 1995). Ketone bodies, namely acetoacetate (AcAc), β-hydroxybutyrate (3-hydroxybutyrate, 3-HB) and acetone, which are overproduced from fatty acids in liver during the conditions of reduced carbohydrate availability such as diabetes and starvation, are released into blood, and serve as a kind of vital alternative metabolic fuel for extrahepatic tissues including brain, skeletal muscle and heart, where they are converted to acetyl-CoA and oxidized in tricarboxylic cycle (TCA) for the provision of large amount of energy (Cahill, 2006; D’Acunzo et al., 2021; Dentin et al., 2006; Krishnakumar et al., 2008; Puchalska and Crawford, 2017; Robinson and Williamson, 1980). Previous studies have shown that the acetate increased significantly in diabetic status and prolonged starvation (Akanji et al., 1989; Seufert et al., 1984; Todesco et al., 1993), and acetate has been considered as a nutrient that nourishes organism by conversion to acetyl-CoA for further catabolism in TCA (Lindsay and Setchell, 1976; Liu et al., 2018; Schug et al., 2015; Schug et al., 2016). Inversely, acetyl-CoA can also be hydrolyzed to acetate by corresponding acyl-CoA thioesterases (ACOTs) family protein (Swarbrick et al., 2014; Tillander et al., 2017). Unfortunately, it is not very clear where, under what condition and how acetate is produced, and what is its biological significance. Considering the high similarity of acetate and ketone body in their production (from acetyl-CoA) and catabolism (converted back to acetyl-CoA), we thoroughly investigated the production and utilization of acetate with ketone bodies as a control and suggest that acetate is an emerging novel “ketone body” that plays important roles similar to classic Ketone Bodies in the energy stresses such as diabetes mellitus and prolonged starvation.

Note: Here our description of acetate as an emerging novel “ketone body” is not aimed to consider it as a real ketone in structure, but to emphasize the high similarity of acetate and the classic Ketone Bodies in the organ (liver) and substrate (fatty acids-derived acetyl-CoA) of their production, the roles they played (as important sources of fuel and energy for many extrahepatic peripheral organs), the feature of their catabolism (converted back to acetyl-CoA and degraded in TCA cycle), as well as the physiological conditions of their production (energy stresses such as prolonged starvation and untreated diabetes mellitus).

## Results

### Acetate is dramatically elevated in energy stresses in mammals

To investigate if acetate is produced as do ketone bodies, we detected the serum glucose, 3-HB, AcAc and other metabolites of 17 diabetes mellitus patients with 8 healthy volunteers as control (**Figure 1—figure supplement 1A; Figure 1—source data 1**). We observed a significant increase of acetate in parallel with the raising of canonical ketone bodies (3-HB and AcAc) and serum glucose in diabetes mellitus patients as compared with healthy control (**Figure 1A**). We then detected acetate in mouse models and found that the levels of serum acetate and ketone bodies were dramatically elevated to the same extent in streptozotocin (STZ)-induced type I diabetic C57BL/6 (**Figure 1B**) and BALB/c mice (**Figure 1—figure supplement 1B**) as well as in type II diabetic db/db mice (**Figure 1—figure supplement 1C**). As expected, starvation also leads to marked descending of serum glucose concentration and ascending of serum acetate and ketone bodies’ level in normal C57BL/6 (**Figure 1C**) and BALB/c mice (**Figure 1—figure supplement 2**). These data demonstrate that serum acetate is boosted to the same extent as the canonical ketone bodies in the energy stresses including diabetes mellitus and starvation. For the sake of simplicity, we designate such acetate hereinafter as energy stress-induced acetate (ES-acetate).

**Figure 1.**
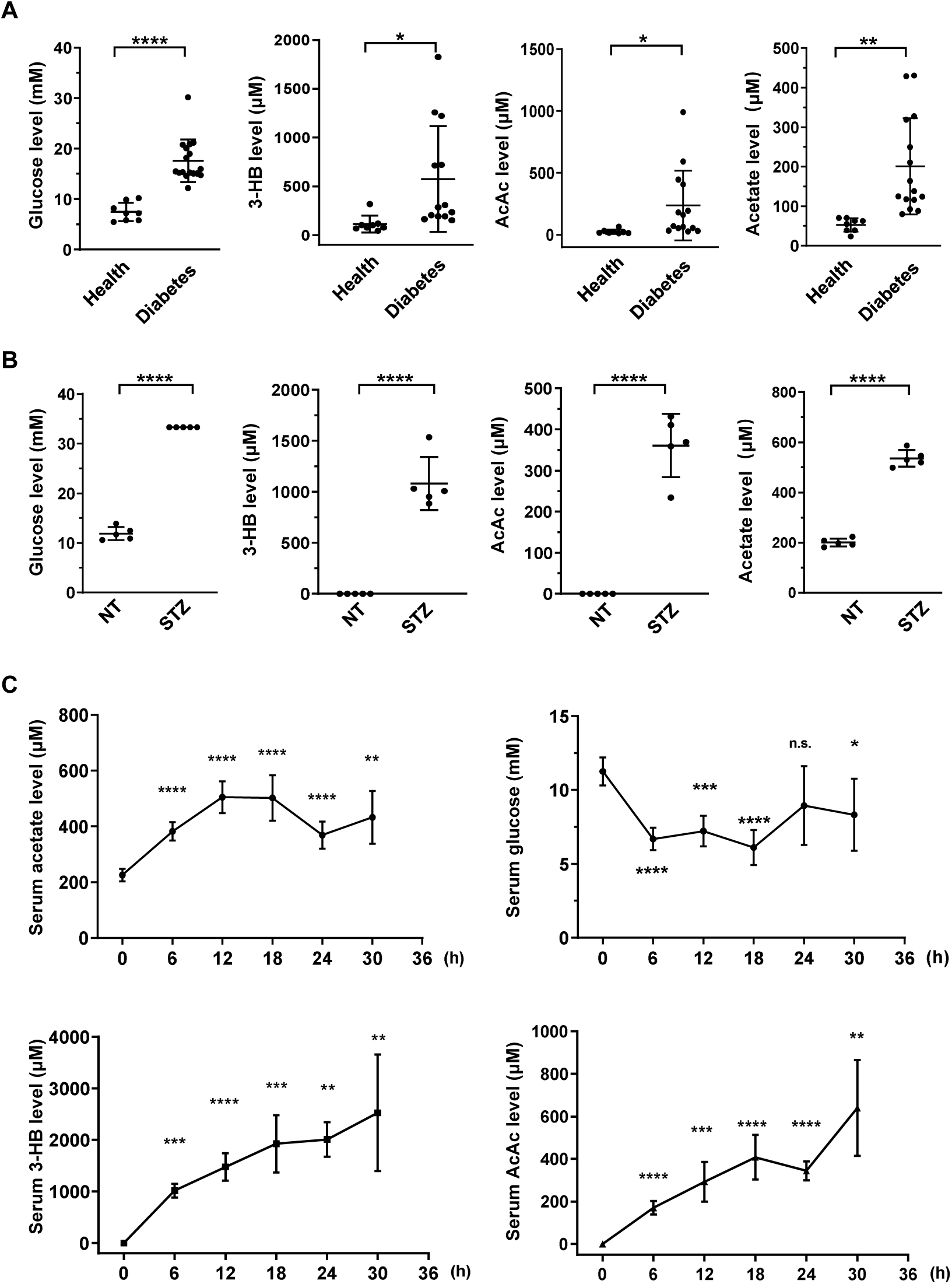
Acetate is produced at a level comparable with ketone bodies in energy stresses. **(A)** Enrichment of glucose, 3-HB, AcAc and acetate in clinical serum samples from healthy volunteers and patients with diabetes mellitus (Health, n=8; Diabetes, n=17). **(B)** Enrichment of glucose, 3-HB, AcAc and acetate in the serum of STZ-induced diabetic mice (C57BL/6, n=5). (**C**) The levels of acetate, 3-HB, AcAc and glucose in the serum of C57BL/6 mice (n=5) starved for indicated time course. Abbreviations: 3-HB, 3-hydroxybutyrate; AcAc, acetoacetate; NT, untreated control; STZ, streptozotocin. Values are expressed as mean±SD and analyzed statistically by two-tailed unpaired Student’s *t* test (\**P*<0.05, \*\**P*<0.01, \*\*\**P*<0.001, \*\*\*\**P*<0.0001, n.s., no significant difference).

### ES-acetate is derived from FFAs in mammalian cells

Next we asked from what nutrients ES-acetate is derived. In mammals, serum acetate contains generally three sources: dietary acetate, metabolic product of gut microbiota and the intermediate of intracellular biochemical processes (Schug et al., 2016). As the mice used above were fed with acetate free diet, we thus focused on gut microbiota and endogenous biochemical reaction. To determine whether gut microbiota contribute to the production of ES-acetate, mice were pre-treated with antibiotics to eliminate gut microbes (saline as control) as reported previously (Sivan et al., 2015). We observed that antibiotics pre-treatment failed to obviously affect the acetate production induced by either starvation (**Figure 2—figure supplement 1A, B**) or diabetes (**Figure 2—figure supplement 1C**), demonstrating that ES-acetate is mainly produced endogenously. Next we detected acetate secreted in culture medium by several cell lines using NMR (**Figure 2—figure supplement 2A, B**) and GC-MS (**Figure 2—figure supplement 2C, D**) and found that these cells showed different ability in producing acetate. Consistently, Liu et al. reported that acetate is derived from glucose in mammalian cells supplied with abundant nutrients (Liu et al., 2018). We observed the secretion of different amounts of U-^13^C-acetate after cells were cultured in medium supplemented with U-^13^C-glucose indeed (**Figure 2—figure supplement 3A**). Interestingly, we also observed the production of 36.6% of non-U-^13^C-acetate, indicating that this proportion of acetate is derived from nutrients other than glucose (**Figure 2—figure supplement 3B**). We then examined if the acetate secreted by cultured cells is derived from amino acids (AAs) and free fatty acids (FFAs) upon starvation. After different cells were cultured in Hanks’ balanced salt solutions (HBSS, free for glucose, fatty acids and amino acids) supplemented with FFAs or amino acids for 20 h, supplementation of FFAs (**Figure 2—figure supplement 4D, E**) rather than amino acids (**Figure 2—figure supplement 4A-C**) significantly increased acetate levels, suggesting the major contribution of FFAs to acetate production. To confirm this observation, a series of widely used cell lines and mouse primary hepatocytes (MPH) were cultured in HBSS supplemented with U-^13^C-palmitate, followed by detection of secreted U-^13^C-acetate (**Figure 2A**). These cell lines displayed quite different ability in conversion of palmitate to acetate and were accordingly divided into FFA-derived acetate-producing cells (FDAPCs: LO_2_, MPH and AML12, etc.) and no-FFA-derived acetate-producing cells (NFDAPCs: HEK-293T and Huh7, etc.). All of the FDAPCs secreted U-^13^C-acetate in a dose-dependent manner of U-^13^C-palmitate supplemented (**Figure 2B-D**). We also observed that high fat diet induced a significant rise of acetate production in both normal and STZ-induced diabetic mice (**Figure 2E**). Taken together, we suggest that acetate can be derived from free fatty acids in the energy stresses.

**Figure 2.**
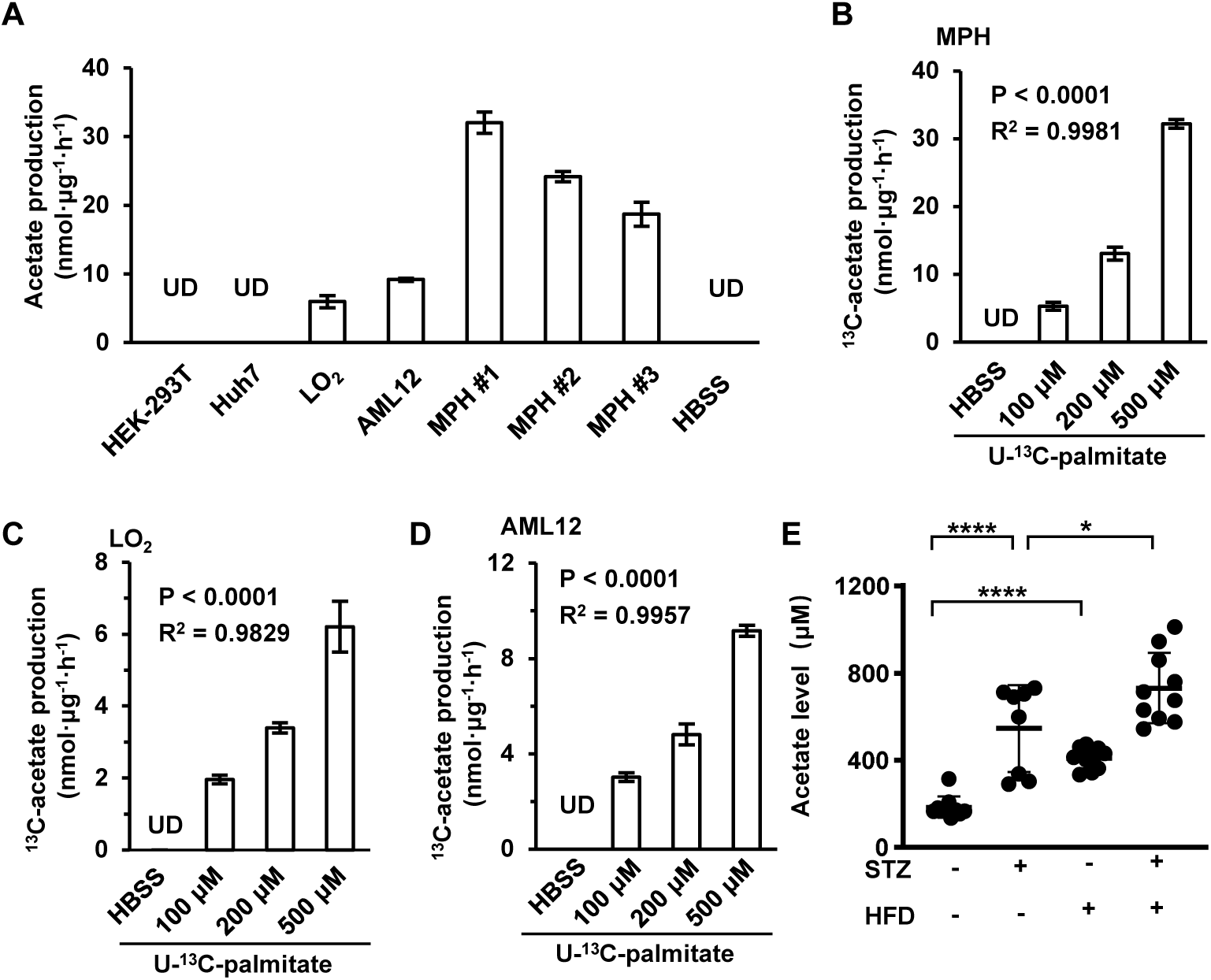
Acetate is derived from FFAs in mammalian cells. (**A**) The amount of U-^13^C-acetate secreted by indicated cells cultured in U-^13^C-palmitate-containing HBSS for 20 h (n=3). (**B**-**D**) The amount of U-^13^C-acetate secreted by MPH (B), LO_2_ (C) and AML12 (D) cells cultured in HBSS supplemented with increasing doses of U-^13^C-palmitate for 20 h (n=3). (**E**) Enrichment of acetate in the serum of untreated or STZ-induced diabetic C57BL/6 mice (n=10) fed with or without high fat diet (HFD). Abbreviations: MPH, mouse primary hepatocytes; UD, undetectable; STZ, streptozotocin. Values are expressed as mean±SD and analyzed statistically by two-tailed unpaired Student’s *t* test (A, E) or one-way ANOVA (B-D), individually (\**P*<0.05, \*\**P*<0.01, \*\*\**P*<0.001, \*\*\*\**P*<0.0001, n.s., no significant difference).

### ACOT12 and ACOT8 are involved in acetate production in mammalian cells

It was reported that acyl-CoAs with different length of carbon chain could be hydrolyzed to FFAs specifically by corresponding acyl-CoA thioesterases (ACOTs) family proteins (Tillander et al., 2017). Acetyl-CoA, the shortest chain of acyl-CoA and the critical product of β-oxidation, is hydrolyzed to acetate by acyl-CoA thioesterase 12 (ACOT12) (Swarbrick et al., 2014). We next analyzed GEO database and found out that the expression of *ACOT1*/*2*/*8*/*12* is upregulated significantly along with the increase of β-oxidation and ketogenesis in mice liver after 24 h of fasting (**Figure 3A**; **Figure 3—figure supplement 1A**). To determine which ACOT is responsible for ES-acetate production, we overexpressed a series of ACOTs in HEK-293T cells and observed large amount of acetate production when either ACOT8 or ACOT12 was overexpressed (**Figure 3B**), indicating the involvement of these two ACOTs in ES-acetate production. Consistently, the protein levels of both ACOT12 and ACOT8 are upregulated robustly in livers of either starved mice or STZ-induced type I diabetic mice (**Figure 3—figure supplement 1B, C**). Furthermore, when ACOT12 and ACOT8 were separately overexpressed in NFDAPCs HEK-293T and Huh7, FFAs-derived acetate was significantly increased (**Figure 3—figure supplement 1D, E**). Similarly, overexpression of wildtype (WT) ACOT12 and ACOT8, rather than their enzyme activity-dead mutants, in HEK-293T (**Figure 3C**) and Huh7 (**Figure 3D**) cells drastically increased U-^13^C-acetate production derived from U-^13^C-palmitate (Ishizuka et al., 2004; Swarbrick et al., 2014). On the contrary, knockdown (KD) of ACOT12 or ACOT8 in FDAPCs MPH (**Figure 3E, F**) and LO_2_ (**Figure 3—figure supplement 1F, G**) diminished U-^13^C-acetate production. These data reveal that ACOT12 and ACOT8 are responsible for ES-acetate production.

**Figure 3.**
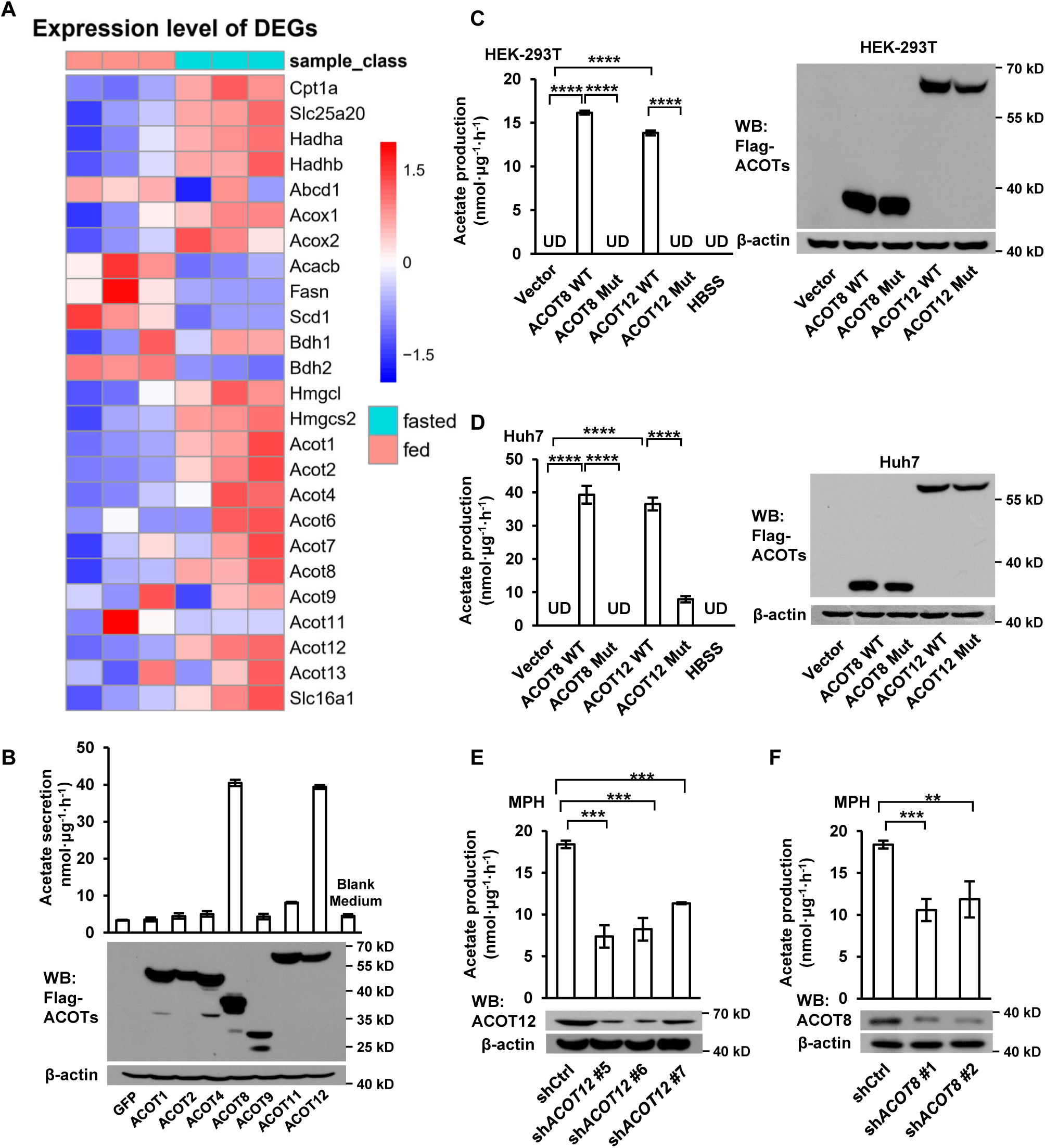
ACOT12 and ACOT8 are involved in acetate production in mammalian cells. (**A**) Heatmap showing hepatic differentially expressed genes between fed group and fasted group, RNAseq analysis data from Goldstein et al. (2017). (**B**) The secretion of acetate (upper panel) by HEK-239T cell lines overexpressing various ACOTs and the protein levels of expressed ACOTs (lower panel). (**C**, **D**) HEK-293T (C) and Huh7 (D) cell lines overexpressing control vector, wildtype (WT) ACOT12 and ACOT8 or their enzyme activity-dead mutants (Mut) were cultured in HBSS containing U-^13^C-palmitate for 20 h, followed by detection of U-^13^C-acetate. (**E**, **F**) U-^13^C-acetate secreted by ACOT12-or ACOT8-knockdown MPH after incubation in U-^13^C-palmitate-containing HBSS for 20 h. Abbreviations: sh*ACOT12*, short hairpin RNA targeting mouse *ACOT12* gene; sh*ACOT8*, short hairpin RNA targeting mouse *ACOT8* gene. UD, undetectable; ACOT8 Mut, ACOT8 H78A mutant; ACOT12 Mut, ACOT12 R312E mutant. Values are expressed as mean±SD (n=3) of three independent experiments and analyzed using unpaired Student’s *t* test (\**P*<0.05, \*\**P*<0.01, \*\*\**P*<0.001, \*\*\*\**P*<0.0001, n.s., no significant difference)

### Hepatic ACOT12 and ACOT8 are responsible for ES-acetate production in energy stresses

Next we were prompted to figure out which organ and subcellular structure are mainly involved in the generation of ES-acetate. Firstly, we analyzed the expression of ACOTs at mRNA level in various tissues of human and mice by employing GTEx and GEO databases, individually. *ACOT12* is mainly expressed in human liver together with ketogenic enzymes (*HMGCS2*, *HMGCSL*, *ACAT1* and *BDH1*), and *ACOT8* is expressed ubiquitously at a relative high level in most tissues (**Figure 4—figure supplement 1**). *ACOT12* is also expressed mainly in mouse liver and kidney, however *ACOT8* seems to be expressed at a much low level in nearly all mouse tissues examined (**Figure 4—figure supplement 2A**). Different from their expression patterns of mRNA in GEO database, we observed high protein levels of both ACOT12 and ACOT8 in mouse liver and kidney (**Figure 4—figure supplement 2B**). Consistently, adenovirus-mediated liver-targeted knockdown of either ACOT12 or ACOT8 dramatically abolished acetate production of starved or diabetic C57BL/6 mice (**Figure 4A-F**) and conditional deletion of ACOT12 or ACOT8 in liver dramatically decreased acetate production in starved mice (**Figure 4G-J**), demonstrating that liver is the main organ responsible for ES-acetate production. Moreover, U-^13^C-acetate derived from U-^13^C-palmitate in glucose free HBSS was diminished by replenishment of glucose (**Figure 4—figure supplement 3**), in accordance with the concept that as energy sources glucose is preferable to fatty acids. These observations demonstrate that hepatic ACOT12 and ACOT8 are induced and responsible for ES-acetate production in diabetes mellitus and during starvation.

**Figure 4.**
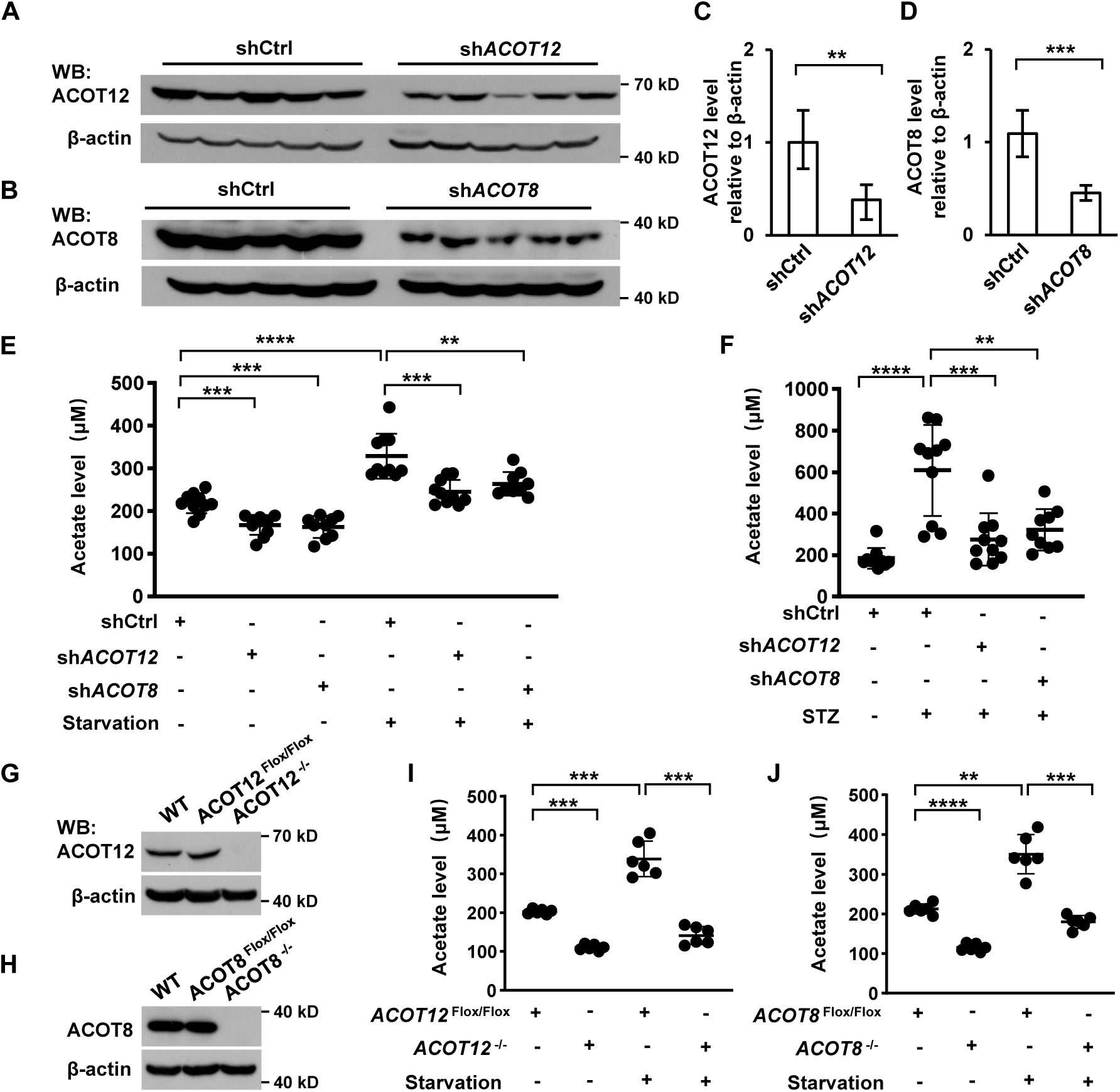
ACOT12 and ACOT8 are responsible for acetate production in energy stresses. (**A**, **C**) ACOT12 in mice (C57BL/6) liver was knocked down by adenovirus-based shRNA, followed by detection of ACOT12 protein with Western Blot (A) and evaluation of knockdown efficiency by calculating ACOT12 level relative to β-actin (C). (**B**, **D**) The knockdown efficiency of ACOT8 was determined as that of ACOT12. (**E**) Enrichment of serum acetate in normal and 16 h fasting mice (C57BL/6) with adenovirus-mediated knockdown of ACOT12 or ACOT8 in liver. (**F**) Enrichment of serum acetate in STZ-induced diabetic mice (C57BL/6) with adenovirus-mediated knockdown of ACOT12 or ACOT8 in liver. (**G**, **H**) ACOT12 (G) or ACOT8 (H) in mice (C57BL/6) liver was conditionally deleted by Cre-Loxp in liver, followed by detection of ACOT12 and ACOT8 protein with Western Blot. (**I**, **J**) Enrichment of serum acetate in normal and 16 h fasting mice (C57BL/6) with Cre-Loxp-mediated conditional deletion of ACOT12 (I) or ACOT8 (J) in liver. Results are expressed as mean±SD of three independent experiments in (C, D), n=10 mice per group in (E, F) and n=6 mice per group in (I, J), and analyzed by using unpaired Student’s *t* test (\**P*<0.05, \*\**P*<0.01, \*\*\**P*<0.001, \*\*\*\**P*<0.0001, n.s., no significant difference).

### ACOT 12&8-catalyzed acetate production is dependent on FFAs oxidation in both mitochondrion and peroxisome

Then we made efforts to clarify in which subcellular domains acetate is produced. Immunofluorescence (IF) staining and cell fractionation showed that ACOT12 was largely localized in cytosol and ACOT8 mainly in peroxisome (**Figure 5A, B**). It’s well-known that fatty acids of different chain length can be oxidized to yield acetyl-CoA in either mitochondria or peroxisome of hepatocyte, and that mitochondrial acetyl-CoA produced in fatty acid oxidation (FAO) is often exported to cytosol in the form of citrate which is further cleaved back to acetyl-CoA by ATP citrate lyase (ACLY) (**Figure 5H**) (Lazarow, 1978; Leighton et al., 1989; Lodhi and Semenkovich, 2014). We thus examined acetate production after mitochondria- or peroxisome-yielded acetyl-CoA had been blocked. Knockdown or etomoxir inhibition of carnitine palmitoyltransferase 1 (CPT1), the main mitochondrial fatty acids transporter, decreased more than one-half of U-^13^C-palmitate-derived U-^13^C-acetate production in LO_2_ cell lines, in spite of mitochondria β-oxidation being nearly completely abolished (**Figure 5C-E**). Similarly, knockdown of ACLY diminished palmitate-derived acetate production to the same extent as CPT1 KD (**Figure 5F**). Then we knocked down ATP binding cassette subfamily D member 1 (ABCD1), a peroxisome fatty acids transporter, and observed less than one-half decline of ^13^C-palmitate-derived U-^13^C-acetate production (**Figure 5G**). These results together with the localization of ACOT12 and ACOT8 suggest that acetyl-CoA produced in FAO of mitochondria and peroxisome is converted to acetate in cytosol by ACOT12 and in peroxisome by ACOT8, individually (**Figure 5H**).

**Figure 5.**
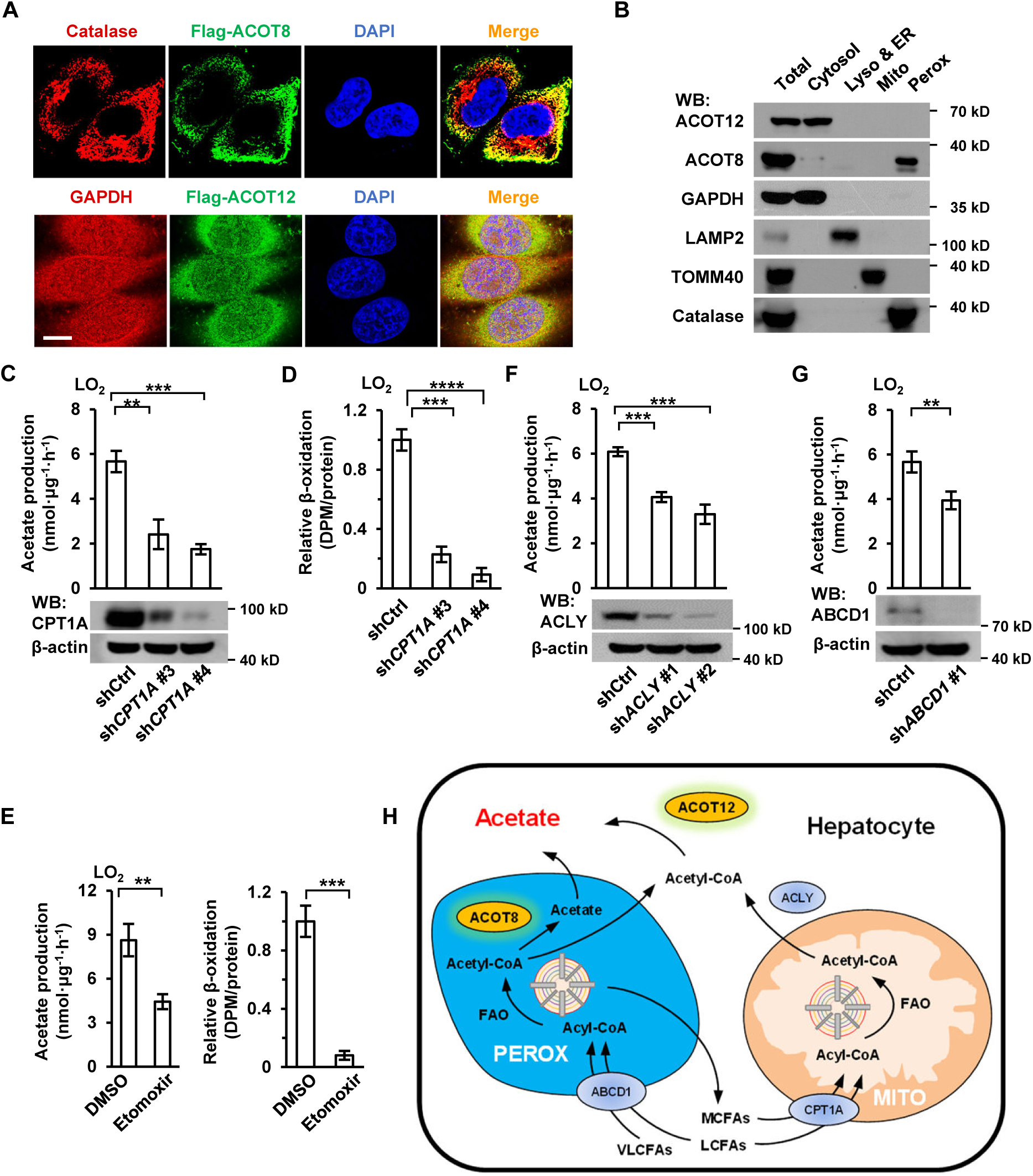
Acetate production is dependent on FFAs oxidation in both mitochondrion and peroxisome. (**A**) Co-immunostaining of Flag-ACOT8 with peroxisome marker catalase and Flag-ACOT12 with cytosol marker GAPDH in LO_2_ cells. Nuclei were stained with DAPI. Scale bars represent 10 μm. (**B**) The protein levels of ACOT12 and ACOT8 in the subcellular fractions of MPH cells. Abbreviations: Lyso, lysosome; ER, endoplasmic reticulum; Mito, mitochondria; Perox, peroxisome. (**C**, **D**) U-^13^C-acetate production (C) and the relative β-oxidation rate (D) in carnitine palmitoyltransferase 1A (CPT1A)-knockdown LO_2_ cells cultured in HBSS containing U-^13^C-palmitate for 20 h. (E) U-^13^C-acetate production (left) and the relative β-oxidation rate (right) of LO_2_ cells cultured in U-^13^C-palmitate-containing HBSS w/wo CPT1 inhibitor etomoxir (20 μM) for 20 h. (**F**) U-^13^C-acetate production in ATP citrate lyase (ACLY)-knockdown LO_2_ cells cultured in HBSS supplemented with U-^13^C-palmitate for 20 h. (**G**) U-^13^C-acetate production in ATP binding cassette subfamily D member 1 (ABCD1)-knockdown LO_2_ cells cultured in HBSS containing U-^13^C-palmitate for 20 h. (**H**) A schematic diagram depicting the mitochondrion and peroxisome pathways of acetate production from FFAs oxidation in hepatocytes. Very long- and long-chain fatty acids (VL/LCFAs) is transported through ABCD1 into peroxisome where it is further degraded into medium-chain fatty acids (MCFAs) via fatty acid oxidation (FAO) process, accompanied by production of acetyl-CoA which is further converted to acetate by peroxisome-localized ACOT8. MCFAs generated in peroxisome are exported into cytosol and absorbed directly by mitochondria. Cytosolic acyl-CoA derived from medium- and long-chain fatty acids (M/LCFAs) is transferred into mitochondria through CPT1A. All fatty acids and acyl-CoA in mitochondria undergo FAO to be degraded to acetyl-CoA. Then acetyl-CoA together with oxaloacetate is synthesized to citrate in TCA cycle, and citrate is exported into cytosol where it is lysed to acetyl-CoA by ACLY. Acetyl-CoA is finally converted to acetate by cytosol-localized ACOT12. Values in (C-G) are expressed as mean±SD (n=3) of three independent measurements. \**P*<0.05, \*\**P*<0.01, \*\*\**P*<0.001, \*\*\*\**P*<0.0001 by two-tailed unpaired Student’s *t* test.

### ACOT12&8-catalyzed recycling of CoA from acetyl-CoA is crucial for sustainable fatty acid oxidation

Afterwards, we tried to explore the biological significance of ES-acetate production in response to energy stresses by detecting a series of serum metabolic parameters. Knockdown of ACOT12 or ACOT8 failed to alter the levels of fasted and non-fasted blood glucose as well as insulin, implying that these two molecules may not be involved in glucose metabolism on non-energy stresses (**Figure 6—figure supplement 1A-C**). However, knockdown of them caused significant accumulation of total FFAs and various saturated or unsaturated fatty acids examined while triacylglycerol (TG) was not altered (**Figure 6—figure supplement 1D-J**). Considering the fact that upon diabetes and prolonged starvation mobilized lipid is mainly transported in the form of plasma albumin-bound fatty acids, rather than TG, these results suggest that ACOT12 and ACOT8 might be required for rapid degradation of fatty acids in these cases. Indeed, we detected attenuated FAO in ACOT12 and ACOT8 knockdown MPH and LO_2_ cells (**Figure 6A, B; Figure 6— figure supplement 1K, L**). Then we were prompted to identify the mechanism underlying such attenuation of FAO. A clue is the knowledge that reduced free Coenzyme A (CoA) is a crucial coenzyme for many metabolic reactions including those involved in oxidative degradation of fatty acid and maintenance of the balance between reduced CoA pool and oxidized CoA pool is definitely important for the sustainment of those reactions (Sivanand et al., 2018). We spontaneously wanted to know if ACOT12&8-catalyzed conversion of acetyl-CoA to free CoA plays a key role in maintaining free CoA level and the balance between reduced CoA and oxidized CoA. To our surprise, in ACOT12/8 KD MPHs, the levels of reduced CoA was decreased by 75.2% and 68.3%, acetyl-CoA increased for 3.49 and 1.71 folds, and the ratios of reduced CoA to acetyl-CoA declined from 7.62 to 0.41 and 0.89, separately (**Figure 6C-E**). In accordance with such alteration, other oxidized CoAs (octanoyl-CoA, caproyl-CoA and succinyl-CoA) whose generation requires sufficient reduced CoA as coenzyme were diminished (**Figure 6—figure supplement 2A-C**). In contrast, metabolites (acetoacetyl-CoA; cholesterol, CHOL; high density lipoprotein cholesterol, HDL-C; low density lipoprotein cholesterol, LDL-C) with acetyl-CoA as the direct substrate for their synthesis were markedly increased (**Figure 6—figure supplement 2D-G**). It is important to point out that among all oxidized CoA examined, the level of acetyl-CoA is far higher than others and the switch between acetyl-CoA and reduced CoA plays a significant role in regulation of CoA pool balance (**Figure 6F**; **Figure 6—figure supplement 2H**). This observation explains why ACOT12- and ACOT8-catalyzed hydrolysis of acetyl-CoA to free CoA and acetate is the crucial step for the maintenance of reduced CoA level and sustained FAO.

**Figure 6.**
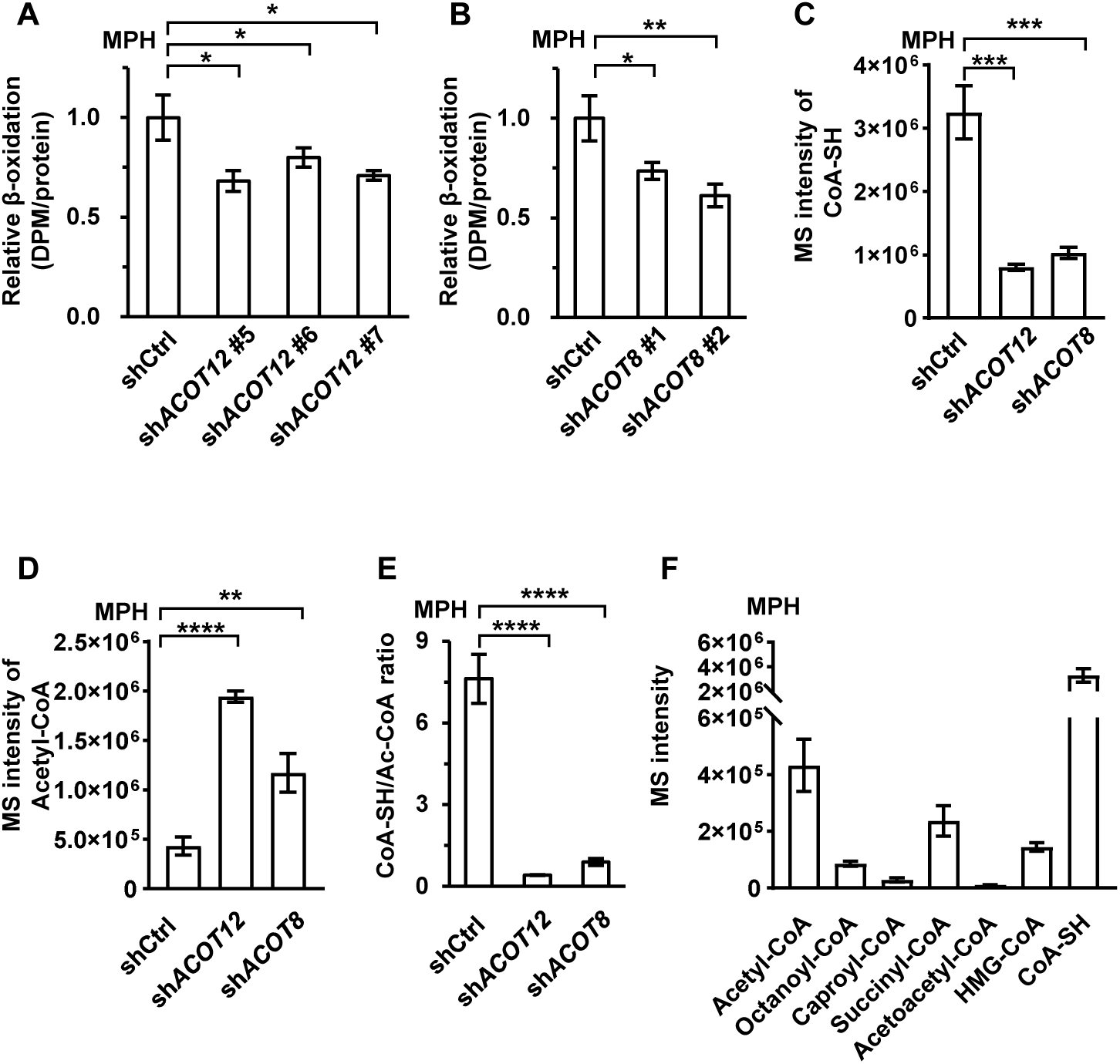
ACOT12 and ACOT8 serve to maintain CoA pool for sustained FAO. (**A**, **B**) MPHs knocked down for ACOT12 (A) or ACOT8 (B) were cultured in glucose free reaction buffer containing 0.8 μCi/mL [9,10-^3^H(N)]-oleic acid for 20 h, followed by determination of the relative β-oxidation rate. (**C**) Relative abundance of reduced CoA in MPHs knocked down for ACOT12 or ACOT8. (**D**) Relative abundance of acetyl-CoA in MPHs knocked down for ACOT12 or ACOT8. (**E**) The ratio of reduced CoA to acetyl-CoA in MPHs knocked down for ACOT12 or ACOT8. (**F**) Relative abundance of reduced CoA and various oxidized CoA in MPHs. Abbreviations: Ac-CoA, acetyl-CoA; HMG-CoA, 3-hydroxy-3-methylglutaryl-CoA. Values are expressed as mean±SD (n=3) of three independent experiments and analyzed using unpaired Student’s *t* test (\**P*<0.05, \*\**P*<0.01, \*\*\**P*<0.001, \*\*\*\**P*<0.0001, n.s., no significant difference).

### Hydrolysis of acetyl-CoA by ACOT12&8 is beneficial to ketogenesis

Distinct from other oxidized CoA, HMG-CoA, a key intermediate for ketone bodies’ synthesis with acetyl-CoA as a substrate, was declined dramatically in ACOT12 and ACOT8 KD MPHs, demonstrating that ACOT12/8 may be positive regulators of HMG-CoA level (**Figure 7A**). Accordingly, the main ketone bodies AcAc and 3-HB were decreased significantly in STZ-induced diabetic mice with knockdown of ACOT12 or ACOT8 (**Figure 7B, C**). To clarify the mechanism underlying ACOT12/8 regulation of HMG-CoA, we detected the protein level of 3-hydroxy-3-methylglutaryl-CoA synthase 2 (HMGCS2), the key enzyme for HMG-CoA synthesis. Interestingly, HMGCS2 was remarkably downregulated in ACOT12/8 KD MPHs (**Figure 7D-G**), indicating that ACOT12&8 are positive regulators of HMGCS2 protein level. A previous study shows that HMGCS2 activity is suppressed by acetylation (Wang et al., 2019). We thus examined the acetylation of HMGCS2 and observed a clear increase of its acetylation in ACOT12/8 KD MPHs (**Figure 7H**), and such alteration is corresponding to the increase of acetyl-CoA level (**Figure 6D**), the direct substrate of acetylation. This observation demonstrates that ACOT12&8 are also positive regulators of HMGCS2 activity by hydrolyzing acetyl-CoA to avoid accumulation of acetyl-CoA and over-acetylation of HMGCS2. Taken together, we suggest that upon energy stress, ACOT12/8 are upregulated and in turn enhance the function of HMGCS2 by increasing not only its amount but also its activity, facilitating ketone body production to fuel the extrahepatic tissues.

**Figure 7.**
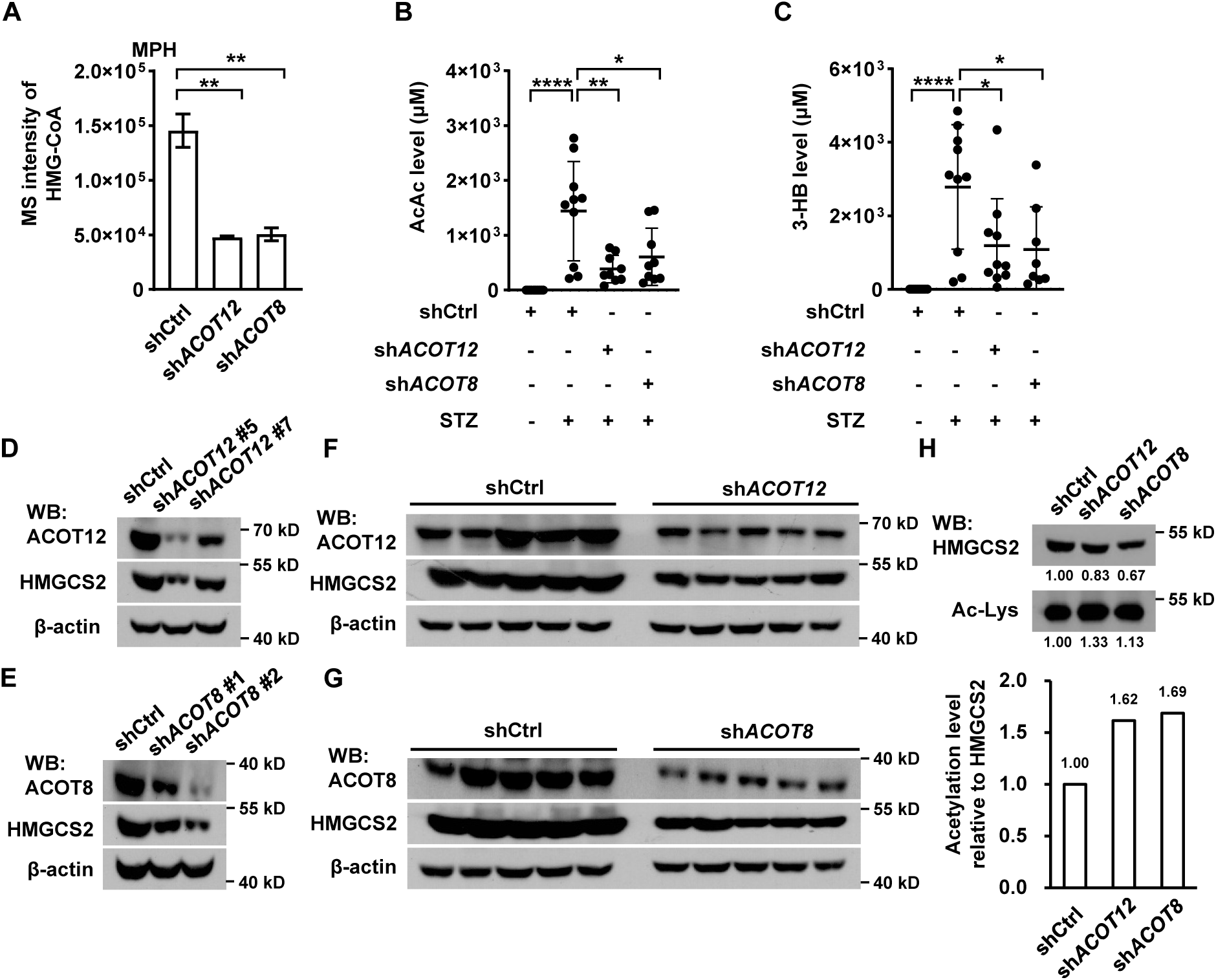
ACOT12 and ACOT8 are required for ketone bodies’ production in STZ-induced diabetes. (**A**) Relative abundance of HMG-CoA in MPHs knocked down for ACOT12 or ACOT8 (n=3). (**B**, **C**) Serum levels of AcAc (B) and 3-HB (C) in STZ-induced diabetic C57BL/6 mice with adenovirus-mediated knockdown of ACOT12 or ACOT8 in liver. (**D**, **E**) The protein levels of HMGCS2 in MPHs knocked down for ACOT12 (D) and ACOT8 (E). (**F**, **G**) ACOT12 (F) and ACOT8 (G) in mice (C57BL/6) liver were knocked down by adenovirus-based shRNA, followed by detection of HMGCS2 protein with Western Blot. (**H**) Western Blot (upper panel) and evaluation of the relative acetylation (Ac-Lys) level by calculating HMGCS2 acetylation relative to HMGCS2 (lower panel). Abbreviations: HMG-CoA, 3-hydroxy-3-methylglutaryl-CoA; HMGCS2, 3-hydroxy-3-methylglutaryl-CoA synthase 2. Results are expressed as mean±SD of three independent experiments in (A) and n=10 mice per group in (B, C), and analyzed by using unpaired Student’s *t* test (\**P*<0.05, \*\**P*<0.01, \*\*\**P*<0.001, \*\*\*\**P*<0.0001, n.s., no significant difference).

### Acetate is beneficial to extrahepatic tissues during energy stresses

It has been well studied that the ketone bodies are produced mainly in liver in diabetes mellitus and prolonged starvation and in turn fuels crucial extrahepatic organs like brain (Puchalska and Crawford, 2017; Robinson and Williamson, 1980). Given acetate was also reported to serve as an energy substance for cells (Comerford et al., 2014; Mashimo et al., 2014; Schug et al., 2016), we spontaneously want to know whether ES-acetate plays the same role as ketone bodies in the same emergency status. 2-^13^C-acetate was injected in starved or STZ-induced diabetic mice intraperitoneally, followed by analysis of ^13^C-labelled metabolic intermediates via LC-MS. ^13^C-acetyl-CoA and ^13^C-incorporated TCA cycle metabolites such as citrate, aconitate, isocitrate, succinate, fumarate and malate were dramatically increased in brain (**Figure 8A-G**), but decreased in muscle (**Figure 8—figure supplement 1**) of starved or diabetic mice as compared with untreated control mice, in line with the notion that brain has priority in energy expenditure during energy stresses. Moreover, we performed intraperitoneal injection of both acetate and 3-HB simultaneously in fasting mice and compared the serum concentration curves of them. Interestingly, acetate took not only less time to reach peak plasma level than 3-HB (5 min vs 12 min), but also much less time to be eliminated (20 min vs 120 min), implying that acetate may be more rapidly absorbed and consumed than 3-HB by extrahepatic organs of mice (**Figure 8H**) as previously reported (Sakakibara et al., 2009). These results suggest that acetate is an emerging novel “ketone body” produced in liver from FFA and functioning to fuel extrahepatic organs, in particular brain, in the emergency status such as energy stresses. Next, we wondered the physiological significance of ES-acetate to animal behaviors under energy stresses and performed a list of behavioral tests: forelimb grip force test for assessing forelimb muscle strength, rotarod test for examining neuromuscular coordination, elevated plus maze test (EPMT) for assessing anxiety-related behavior, Y-maze test (YMZT) and novel object recognition (NOR) test for evaluating working memory and cognitive functions. It’s clear that the forelimb strength and running time in rotarod test were dramatically declined in diabetic mice, further deteriorated by knockdown of ACOT12 or ACOT8, and rescued by administration of exogenous acetate (**Figure 8—figure supplement 2A, B**). Interestingly, the parameters related to muscle force and movement ability in other tests including total distance in YMZT, total entries in YMZT and total distance in NOR test were also decreased in diabetic mice and further worsened by knockdown of ACOT12 or ACOT8 (**Figure 8—figure supplement 2E, F and H**). These observations demonstrate that ES-acetate is important for muscle force and neuromuscular coordinated movement ability. EPMT test, correct alteration in YMZT and object recognition index in NOR test showed no significant difference among normal mice, diabetic mice and ACOT12/8 KD mice (**Figure 8—figure supplement 2C, D and G**), indicating that psychiatric, memory and cognitive behaviors is not markedly influenced by ACOT12/8 KD in early stage of diabetes mellitus, possibly due to brain exhibits the highest flexibility in utilizing all kinds of available energy sources such as glucose, ketone body and acetate in energy stresses among all extrahepatic tissues.

**Figure 8.**
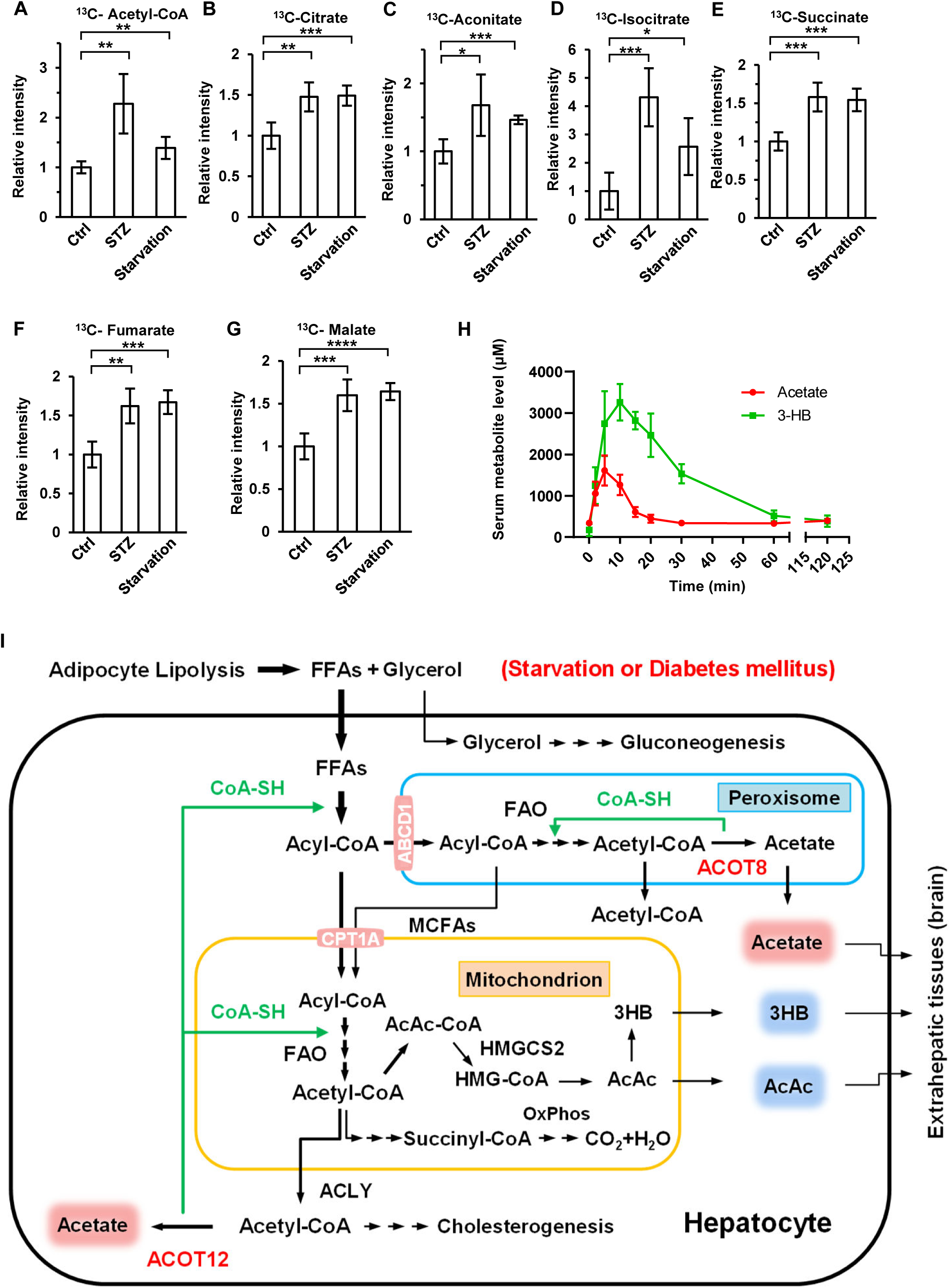
Brain exhibits increased acetate consumption during energy stresses. (**A**-**G**) Relative abundance of ^13^C-acetyl-CoA (A), ^13^C-citrate (B), ^13^C-aconitate (C), ^13^C-isocitrate (D), ^13^C-succinate (E), ^13^C-fumarate (F) and ^13^C-malate (G) in the brain of starved or diabetic mice (C57BL/6) was determined 1 h after intraperitoneal injection of 2-^13^C-acetate (310 mg/kg). (**H**) The abundance of acetate and 3-HB in the serum of fasting mice (C57BL/6) after intraperitoneal injection (acetate 300 mg/kg, 3-HB 520 mg/kg). (**I**) A working model describing the biological significance of ACOT12- and ACOT8-catalized conversion of acetyl-CoA to acetate and CoA. In the status of energy stress such as diabetes mellitus and prolonged starvation, body takes at least two advantages by converting acetyl-CoA to acetate and CoA: 1) CoA is required for sustained FAO and ketone bodies production in liver; 2) acetate serves as a novel ketone body to fuel extrahepatic tissues, particularly brain. Values in (A-H) are expressed as mean±SD (n=5 mice per group in (A-G) and n=7 mice per group in (H)) and analyzed statistically by employing unpaired Student’s *t* test (\**P*<0.05, \*\**P*<0.01, \*\*\**P*<0.001, \*\*\*\**P*<0.0001, n.s., no significant difference).

## Discussion

It is well known that glucose and ketone bodies are the main fuels for brain in different physiological conditions. In normal condition brain uses glucose as the main energy source. In contrast, it utilizes ketone bodies as an important energy source in the status of energy stress such as diabetes mellitus and prolonged starvation, because in these cases body glucose storage has been already exhausted and gluconeogenesis cannot provide sufficient glucose. As a result, sustained mobilization of stored lipid and the resultant production of ketone bodies from fatty acid oxidation is crucial for nourishing extrahepatic tissues, in particular brain in energy stresses. However, sustained fatty acid oxidation in hepatocyte needs rapid recycling of free CoA which is the crucial co-enzyme for FAO. Our study elucidates that such requirement is satisfied by ACOT12- and ACOT8-catalized conversion of acetyl-CoA to acetate and CoA. It is important to point out that the significance of ACOT12- and ACOT8-catalized recycling of free CoA for FAO is analogous to that of lactate dehydrogenase (LDH)-catalyzed recycling of NAD^+^ for glycolysis (Castro et al., 2009; Cerdan et al., 2006). In addition to providing CoA, this reaction also provide acetate which serves as an energy source to fuel extrahepatic tissues (**Figure 8I**). As an alternative fuel in emergency status, acetate may be preferred by brain, because it is directly converted to acetyl-CoA by acetyl-CoA synthetase (ACSS), undoubtedly more convenient than acetoacetate and 3-HB which need 2 and 3 enzyme-catalyzed steps to be converted to acetyl-CoA, individually. In this regard, we suggest to consider acetate as an emerging novel “ketone body” and its blood level should be detected, together with classic Ketone Bodies, to indicate the status of FAO in liver and lipid mobilization in adipose tissue upon energy stress.

In summary, in this study we clarify where and how acetate is produced in energy stresses and identify the profound biological significance of acetate production catalyzed by ACOT 12&8. More importantly, we suggest that acetate is an emerging novel “ketone body” and may be used as a parameter to evaluate the progression of energy stress in the future.

## Materials and methods

### Collection of clinical samples

Human clinical serum samples were collected based on ethical approval of the clinical research ethics committee of the First Affiliated Hospital of Xiamen University (Xiamen, Fujian, China). The information of patients (diagnosed with type II diabetes) and healthy volunteers is provided in **Figure 1—source data 1**. The serum samples were stored in -80 °C refrigerator and mainly obtained from The First Affiliated Hospital of Xiamen University (China) after obtaining informed consent.

### Animal studies

All animal studies were approved by the Animal Ethics Committee of Xiamen University (China) (acceptance no: XMULAC20190166). BALB/c and C57BL/6 mice (6-7 weeks, random sex, in groups) were obtained from Xiamen University Laboratory Animal Center (China). C57BLKS/J-LepR^db^/LepR^db^ (db/db) mice were purchased from GemPharmatech Co, Ltd (China). All Animals were kept in SPF condition with 12 h light-dark cycle, free chow and water accessed to standard rodent diet in accordance with institutional guidelines. For STZ induced diabetic models (Gonzalez et al., 2003; Like and Rossini, 1976), mice were randomized and fasted for 12 h but water is allowed before intraperitoneal injection of STZ (a single high dose of 150 mg/kg). Note that BALB/c diabetic mice induced by STZ need to be fed with a 60 kcal% fat diet (high fat diet, HFD) for acetate detection until the end of experiment. For animal starvation experiment, mice were fasted but water is allowed. For antibiotic treatment experiment, mice were treated with a mixture of antibiotics including 1 mg/ml ampicillin, 5 mg/ml streptomycin and 1 mg/ml colistin in sterile drinking water for 3 weeks before STZ injection or starvation experiment and treatment was continued to the end of experiment (Vetizou et al., 2015). For Cre-Loxp-mediated liver-specific ACOT12 or ACOT8 knockout mice (*Acot12*^-/-^ or *Acot8*^-/-^), C57BL/6JGpt-*Acot12*^em1Cflox^/Gpt and C57BL/6JGpt-*Acot8*^em1Cflox^/Gpt mice were purchased from GemPharmatech Co, Ltd (China), and *Acot12* ^Flox/Flox^ or *Acot8* ^Flox/Flox^ mice cross with *Alb-Cre* mice (C57BL/6), confirm efficient deletion of ACOT12 or ACOT8 specifically in liver at protein (6-7 weeks, random sex, in groups). For adenovirus-mediated liver-specific RNAi mouse models, injection of 200 μL adenovirus (titer: 10^12^) via tail vein was performed in C57BL/6 mice (6-7 weeks, random sex, in groups) and knockdown efficiency in liver was determined by Western Blot after 2 months of injection. Adenoviruses were propagated in QBI-293A cells and purified by cesium chloride density gradient ultracentrifugation. For all animal models, blood was collected from tail vein, followed by detection of glucose with Roche glucometer and other serum components with NMR and MS. Tissue samples of mice were collected at the end of experiment for Western Blot.

### Animal behavior analysis

Behavior analysis were performed by using age-matched C57BL/6 mice (4 months, random sex, in groups). Diabetes mellitus, as mentioned above, was induced by STZ in normal or adenovirus-mediated liver-specific RNAi mice. Mice were moved to the experimental room 1 h before starting the experiment. All objects or apparatus were thoroughly cleaned with 75% alcohol between trials to remove odors. And all mice exhibited excellent health throughout the study period. For Forelimb Grip Force Test, the forelimb grip forces were measured by a Grip Strength Meter (Ugo Basile, Italy) and the peak force was defined as the average of three successive measurements. The rotarod test was performed by a progressive acceleration setting from 5 to 40 rpm for 2 min using a five-lane apparatus (Ugo Basile, Italy). Before rotarod test, all mice were trained under a condition of 5 rpm 2 min daily for 2 days. For acetate rescuing experiments, the forelimb grip force test and rotarod test were conducted after 5 min of intraperitoneal injection of acetate. The Elevated Plus Maze Test (EPMT) was performed with an elevated plus maze (40 cm in length, 10 cm in width, 50 cm in height, Panlad, Spain) which consists of four elevated arms radiating from a central platform, forming a plus shape. Two of the opposed arms were enclosed by a wall of 20 cm in height. Each mouse was placed in the same area, and then left to explore the maze for 5 min. The amount of time spent in the open and closed arms was measured with a video-imaging system (Dazzle DVC100 Video). Data analyses were performed using active-monitoring software (smart3.0). Y-Maze Test (YMZT) was carried out using a Y-shaped maze with three light-colored, opaque arms (30 cm in length, 6 cm in width, 5 cm in height, Panlad, Spain) orientated at 120 angles from each other. Each mouse was placed in the same area, and then left to explore the maze for 5 min. The number of entries into the arms and alterations were recorded with a video-imaging system (Dazzle DVC100 Video). Data analyses were performed using active-monitoring software (smart3.0). The one-trial Novel Object Recognition (NOR) Test was carried out using an open-field apparatus (40×40×40 cm, Panlad, Spain) as test box and the protocol consisted of two test sessions separated by an over 20-min delay during which mice were returned to their home cage. In each test session, every mouse was placed in the same area, and then left to explore the open field for 5 min. For the first session, the mice were trained in the arena where two cubes (5 × 5 × 5 cm, familiar object) were placed as objects A and B. For second session, the mice were trained in the arena where one object A and one cylinder (5 cm diameter, 5 cm height, novel object) designated as object C were placed. The time and distance of novel and familiar objects exploration were recorded with a video-imaging system (Dazzle DVC100 Video) during the trials of each session. Data analyses were performed using active-monitoring software (smart3.0). The ratio of object C exploration time to total time represents the object recognition index.

### Plasmids constructs

Full-length cDNAs encoding human ACOTs (gene ID: 25082 for ACOT1, gene ID: 15824 for ACOT2, gene ID: 9637 for ACOT4, gene ID: 24012 for ACOT8, gene ID: 17595 for ACOT9, gene ID: 10617 for ACOT11 and gene ID: 134526 for ACOT12) was obtained from Core Facility of Biomedical Sciences, Xiamen University. Point mutation of ACOT12 (ACOT12 R312E&R313E) (Lu et al., 2019) and ACOT8 (ACOT8 H78A) (Ishizuka et al., 2004) were constructed by PCR-mediated mutagenesis using PrimerSTAR DNA polymerase (Takara). cDNAs for proteins expression were constructed in pLV cs2.0 vectors. shRNAs were constructed in lentivirus-based pLL3.7 vector. shRNAs against mouse *ACOT12* (#5, #6 and #7) and mouse *ACOT8* (#1 and #2) were also constructed in pAdEasy-1 (Stratagene) for adenovirus packaging based on The AdEasy^TM^ Technology (He et al., 1998).

### Cell culture, transfections and cell treatments

HeLa, HEK-293T, HT1080, Huh7, LO_2_, H3255, A549, QBI-293A and HEB cell lines were taken from our laboratory cells bank and authenticated by Short Tandem Repeat (STR) profiling analysis. HCT116, 786-O, HepG2, Hepa1-6 and AML12 cell lines were obtained from and authenticated by Cell Bank of the Chinese Academy of Sciences (Shanghai). All cells were examined negative for mycoplasma infection using PCR-based Mycoplasma Detection Kit (Sigma, MP0035-1KT). Mouse embryonic fibroblast cell (MEF) was isolated from embryos of mice at 13.5 days post-coitum and further immortalized by infection of the SV-40 larger T antigen expressing retroviruses. All the cell lines were cultured in DMEM (Gibco) with 10% fetal bovine serum (FBS, Gemini) at 37 °C in incubator containing 5% CO_2_. HEK-293T was used for transient transfection and lentivirus package with polyethylenimine (PEI, 10 μM, Polyscience) as transfection reagent. The virus-containing medium was collected after 24 hours of transfection, filtered by 0.45 μm Steriflip filter (Millipore) and stored at -80 °C for infection. The infected cells were passaged until stable cell lines were constructed. For all kinds of treatment, cells were seeded in 35 mm dishes and cultured for 24 h before treatment. To measure glucose-derived acetate, cells were rinsed with PBS and then incubated in glucose-free DMEM (1 mL, Gibco) supplemented with 10% FBS and 10 mM U-^13^C-glucose for 20 h before harvest. To measure fatty acid-derived acetate, cells were rinsed with PBS and then incubated in Hanks’ balanced salt solution (HBSS) supplemented with 10% FBS and 500 μM bovine serum albumin (BSA, fatty acids free, Yeasen Biotech)-conjugated free fatty acid (Myristate, Palmitate, Stearate or U-^13^C-palmitate) for 20 h before harvest. To determine amino acid-derived acetate, cells were rinsed with PBS and then incubated in HBSS supplemented with 10% FBS and 2× or 4× amino acids for 20 h before harvest. 2× amino acids contains double concentrations of amino acids in DMEM and 4× contains quadruple concentrations of amino acids in DMEM. HBSS media (1 L, pH 7.4) contains CaCl_2_ (140 mg), MgCl_2_·6H_2_O (100 mg), MgSO_4_·7H_2_O (100 mg), KCl (400 mg), KH_2_PO_4_ (60 mg), NaHCO_3_ (350 mg), NaCl (8 g) and Na_2_HPO_4_ (48 mg). To determine gluconeogenesis, cells were rinsed with PBS and then incubated in HBSS media (glucose-free) supplemented with 100 nM glucagon (Acmec, G78830) for 4 h before harvest as previously described (Liu et al., 2017).

### Adenovirus packaging and infection

Sterile linearized recombinant AdEasy^TM^ plasmids were transfected in QBI-293A cell lines with Turbofect transfection reagent for adenovirus packaging as described previously (He et al., 1998). Fresh QBI-293A cells were further infected by the primary adenoviruses for amplification and purification of recombinant adenovirus. The purified adenovirus was used for infection of mouse primary hepatocytes (MPH) in vitro and liver cells in vivo.

### Mouse primary hepatocytes isolation

Mouse primary hepatocytes were obtained from C57BL/6 mice by perfusing the liver through the portal vein with calcium-free buffer A (1 mM EGTA and Kreb-Ringer buffer), followed by perfusion with buffer B (collagenase-IV from Sigma, Kreb-Ringer buffer and 5 mM CaCl_2_). Hepatic parenchymal cells were maintained in DMEM containing 10% FBS and precipitated by centrifugation (50 g for 3 min). Next, the isolated cells were plated in dishes pre-treated by collagen-I (CORNING) and cultured in DMEM with 10% FBS at 37 °C in humid incubator containing 5% CO_2_. Kreb-Ringer buffer (1 L, pH 7.4) contains NaCl (7 g), NaHCO_3_ (3 g), HEPES (5 mM, pH 7.45), Solution C (10 ml) and Glucose (1 g). Solution C contains KCl (480 mM), MgSO_4_ (120 mM) and KH_2_PO_4_ (120 mM). All buffers and media above contain penicillin (100 IU, Sangon Biotech) and streptomycin (100 mg/ml, Sangon Biotech) (Huang et al., 2011).

### Immunoprecipitation and Western Blot

Cells or tissues were harvested in a lysis buffer (20 mM Tris-HCl (pH 7.4), 150 mM NaCl, 1 mM EDTA, 2.5 mM sodium pyrophosphate, 1 mM β-glycerolphosphate, 1 mM sodium orthovanadate, 1 mM EGTA, 1% Triton, 1 μg/ml leupeptin, 1 mM phenylmethylsulfonyl fluoride), sonicated and centrifuged at 20,000 g for 15 min at 4°C. For immunoprecipitation, incubated the supernatant with corresponding antibody for 12h at 4°C and then incubated with A/G plus-agarose beads (Santa Cruz Biotechnology, Inc.) for 2h at 4°C. For Western Blot, immunoprecipitates or total cell lysates supernatant added with SDS loading buffer, boiled for 10 min and separated by SDS-PAGE, followed by transferring to PVDF membranes (Roche). The PVDF membranes were incubated with specific antibodies for 3 h and proteins were visualized by enhanced chemiluminescence (ECL) system. The intensity of blots was analyzed by using Image J.

### Subcellular fraction purification

For subcellular fraction purification, the Peroxisome Isolation kit (Sigma) was used to isolate peroxisomes from primary hepatocytes by referring to the protocol provided by Sigma-Aldrich. The isolated subcellular fractions were lysed with lysis buffer and analyzed by Western Blot.

### Immunofluorescence

LO_2_ cells grown on coverslips at 30%-40% of confluence were washed with PBS and fixed in 4% paraformaldehyde for 10 min. The fixed cells were treated with 0.2% Triton X-100 in PBS for 10 min at room temperature to permeabilize membrane and then incubated with 5% BSA in TBST (20 mM Tris, 150 mM NaCl, and 0.1% Tween 20) for 1h to block non-specific binding sites. Next cells were incubated with primary antibodies diluted in TBST containing 5% BSA for 1 h at room temperature and washed three times with 0.02% Triton-X100 in PBS, followed by incubation with fluorescent secondary antibodies for 1h. After washed three times with 0.02% Triton-X100 in PBS, all coverslips were counterstained with DAPI and mounted on microscope slides with 90% glycerol. Images were captured by Leica TCS SP8 confocal microscope at pixels of 1024×1024.

### Biochemical analyses

To prepare mouse serum, mouse blood was collected into 1.5 mL Eppendorf tube and allowed to clot for 30 min at 4°C. Then samples were centrifuged for 30 min (1300 g) at 4°C and the serum layer was carefully moved into a new 1.5 mL Eppendorf tube. Plasma levels of TG, CHOL, HDL-C and LDL-C were measured in Clinical Laboratory of Zhongshan Hospital, affiliated to Xiamen University. Plasma insulin levels were measured using the MOUSE INS-1055 ELISA KIT (Meikebio) with a standard curve, following the manufacturer’s protocol. Plasma total FFAs levels were measured using the free fatty acid (FFA) content assay kit (Beijing Boxbio Science & Technology) with a standard curve, following the manufacturer’s protocol and each FFA was measured by gas chromatography mass spectrometry. Fasted, mice were fasted for 12 h; Non-fasted, mice were fed normally.

### Gas Chromatography Mass Spectrometry

To identify the acetate produced by cells, metabolites in culture medium were subjected to acidification and extraction, followed by analysis using gas chromatography mass spectrometry (GC-MS) as previously described with some optimization (Fellows et al., 2018). First, equal propionic acid and butyric acid were added to cell cultured media (100 μL) as internal reference in Eppendorf tubes. Subsequently, 40 mg of sodium chloride, 20 mg of citric acid and 40 μL of 1 M hydrochloric acid were added to acidize metabolites. After acidification, acetate, propionic acid and butyric acid were liable to be extracted by 200 μL of n-butanol. Next, the tubes were vortexed for 3 min and centrifuged at 20,000 g for 20 min. The supernatant was transferred to HPLC vial and 1 μL mixture was determined.

Mouse serum was extracted as mentioned above and subjected to measurement of free fatty acids (FFAs) levels employing GC-MS. In brief, 30 μL of cold mouse serum was transferred to new 1.5 mL Eppendorf tubes and 500 μL of cold 50% methanol (containing 2.5 μg/mL tridecanoic acid as internal reference) was added to the samples, followed by addition of 500 μL of cold chloroform. Next, samples were vortexed at 4 °C for 10 min and centrifuged (12000 g) at 4 °C for 20 min to separate the phase. The chloroform phase containing the total fatty acid content was separated and lyophilized by nitrogen. Dried fatty acid samples were esterified with 100 μL 1% sulfuric acid in methanol for 60 min at 80 °C and extracted by addition of 100 μL n-hexane. The supernatant was transferred to HPLC vial and 1 μL mixture was determined.

Analysis was performed using an Agilent 7890B gas chromatography system coupled to an Agilent 5977B mass spectrometric detector (MSD) and a fused-silica capillary DB-FFAP with dimensions of 30 m×0.25 mm internal diameter (i.d.) coated with a 0.25 µm thick layer. The initial oven temperature was 50 °C, then ramped to 110 °C at a rate of 15 °C min^−1^, to 180 °C at a rate of 5 °C min^−1^, to 240 °C at a rate of 15 °C min^−1^, and finally held at 240 °C for 10 min. Helium was used as a carrier gas at a constant flow rate of 1 mL min^−1^ through the column. The temperatures of the front inlet, transfer line, and electron impact (EI) ion source were set at 240 °C, 260 °C, and 230 °C, respectively. The electron energy was −70 eV, and the mass spectral data was collected in a full scan mode (m/z 30 - 300).

### Liquid chromatography-mass spectrometer

The metabolites of TCA cycle were determined by Liquid chromatography-mass spectrometer (LC-MS) as described (Hui et al., 2020). To prepare the samples for measurement of metabolites in tissues, the equivalent tissues of brain or muscle were quenched by pre-cold methanol solution (methanol: ddH_2_O=4:1) and homogenated, followed by centrifugation (12,000 g, 20 min). The supernatants were collected in new Eppendorf tubes and dried at 4 °C and the pellets were resuspended in acetonitrile solution (acetonitrile: ddH_2_O=1:1) and transferred to HPLC vial. To prepare the samples for measurement of intracellular metabolites of in vitro cultured cells, the medium was discarded and cells cultured in 35 mm dish were gently washed twice by cold PBS, followed by the addition of pre-cold methanol solution (methanol: ddH_2_O=4:1, containing 160 ng/mL U-^13^C-glutamine as internal reference) to each well. Samples were then handled as described above for measurement of tissue metabolites. For analysis of metabolites in above prepared samples, the liquid chromatography with SCIEX ExionLC AD was prepared and all chromatographic separations were performed with a Millipore ZIC-pHILIC column (5 μm, 2.1×100 mm internal dimensions, PN: 1.50462.0001). The column was maintained at 40°C and the injection volume of all samples was 2 μL. The mobile phase that consisted of 15 mM ammonium acetate and 3 ml/L Ammonium Hydroxide (> 28%) in LC-MS grade water (mobile phase A) and LC−MS grade 90% (v/v) acetonitrile in HPLC water (mobile phase B) ran at a flow rate of 0.2 mL/min. The ingredients were separated with the following gradient program: 95% B for 2 min, then changed to 45% B within 13 min (linear gradient) and maintained for 3 min, then changed to 95% B directly and maintained for 4 min. The flow rate was 0.2 mL/min. The QTRAP mass spectrometer used an Turbo V ion source. The ion source was run in negative mode with a spray voltage of -4,500 V, with Gas1 40 psi, Gas2 50 psi and Curtain gas 35 psi. Metabolites were measured using the multiple reactions monitoring mode (MRM). The relative amounts of metabolites were analyzed by MultiQuant Software Software (AB SCIEX).

### NMR measurements

To prepare the culture medium samples for NMR analysis, medium harvested after treatment of cells were centrifuged (12000 g at 4 °C for 10 min) and the supernatants (400 μL) were transferred into 5 mm NMR tubes for NMR measurement. The clinical serum samples (200 μL) were thawed on ice, mixed with 200 μL NMR buffer (50 mM sodium phosphate buffer, pH 7.4 in D_2_O) and centrifuged (12000 g) at 4 °C for 10 min. The supernatants (400 μL) were transferred into 5 mm NMR tubes for NMR measurement. For preparation of mice blood sample, 25 μL blood was mixed with 75 μL saline immediately, and centrifuged (3000 g) at 4 °C for 10 min. The supernatants (100 μL) were mixed with 300 μL NMR buffer and transferred into 5 mm NMR tubes for NMR measurement. An internal-tube containing 200 μL D_2_O (used for field-frequency lock) with 1 mM sodium 3-(trimethylsilyl) propionate-2,2,3,3-d4 (TSP) was used to provide the chemical shift reference (δ 0.00) and quantify the metabolites. NMR measurements were performed on a Bruker Avance III 850 MHz spectrometer (Bruker BioSpin, Germany) equipped with a TCI cryoprobe at 25 °C provided by College of Chemistry and Chemical Engineering (Xiamen University) and a Bruker Avance III 600 MHz spectrometer (Bruker BioSpin, Germany) provided by Core Facility of Biomedical Sciences (Xiamen University). One dimensional (1D) CPMG spectra were acquired using the pulse sequence [RD-90°-(τ-180°-τ) _n_ -ACQ] with water suppression for culture medium and serum samples. For the purpose of metabolite resonance assignments, two dimensional (2D) ^1^H-^13^C heteronuclear single quantum coherence (HSQC) spectra were recorded on selected NMR samples. Identified metabolites were confirmed by a combination of 2D NMR data and the Human Metabolome Data Base (HMDB).

### Fatty acid oxidation measurement

Fatty acid oxidation was carried out as previous described (Li et al., 2018). Briefly, cells were cultured in 35 mm dish and rinsed twice with PBS to remove the residue medium. 1 mL reaction buffer (119 mM NaCl, 10 mM HEPES (pH 7.4), 5 mM KCl, 2.6 mM MgSO_4_, 25 mM NaHCO_3_, 2.6 mM KH_2_PO_4_, 2 mM CaCl_2_, 1 mM BSA-congregated oleic acid and 0.8 μCi/mL [9,10-^3^H(N)]-oleic acid) was added in dish to incubate cells at 37 °C for 12 h, followed by centrifugation (1000 g, 5 min) to obtain supernatant. Then, 192 μL of 1.3 M perchloric acid was added to 480 μL of supernatant. The mixture was centrifuged (20,000 g, 5 min) and 240 μL supernatant was mixed with 2.4 mL of scintillation liquid and 3H radioactivity, followed by measurement with liquid scintillation counter (Tri-Carb 2008TR, Perkins Elmer, USA), provided by Center of Major Equipment and Technology (COMET), State Key Laboratory of Marine Environmental Science, Xiamen University.

### Database analysis

The data of GSE72086 (Goldstein et al., 2017) by RNA-Seq was downloaded from the public NCBI Gene Expression Omnibus (www.ncbi.nlm.nih.gov/geo/), analyzed by limma package (Ritchie et al., 2015) and visualized by ggplot2 and ggrepel packages in R (version 3.6.3). GSE72086 contains 6 samples: 3 fed treatments and 3 fasted-24h treatments. Here, the probe with the greatest *p* value was chosen to determine the differential gene expression for multiple probes corresponding to the same gene. Adjusted *p* value <0.05 and |log fold change (log FC) |≥1 were chosen as the threshold value.

The tissue-specific mRNA expression of target gene was analyzed by using GTEx database (www.gtexportal.org) for the human data and GSE24207 of GEO database for the mice data as reported (Fagerberg et al., 2014; Thorrez et al., 2011).

### Statistical analysis

The two-tailed Student’s *t* test was used to analyze difference between two groups with Graphpad Prism 8 and Excel. One-way ANOVA was used to compare values among more than two groups with Graphpad Prism 9 and R (version 3.6.3). Difference was considered significant if *p* value was lower than 0.05 (\**P*<0.05; \*\**P*<0.01; \*\*\**P*<0.001; \*\*\*\**P*<0.0001).

## Supporting information

Author response

Key Resources Table

MDAR_checklist

## Acknowledgments

We thank members of the Qinxi Li’s laboratory for productive discussions and comments on this manuscript. We thank Center of Major Equipment and Technology (COMET), State Key Laboratory of Marine Environmental Science (Xiamen University) for technical support.

## The List of Figures and Source Data

**Figure 1—source data 1**

The Excel spreadsheet provided contains the source data pertaining to the patient information of clinical data depicted in Figure 1.

**Figure 1—figure supplement 1.**
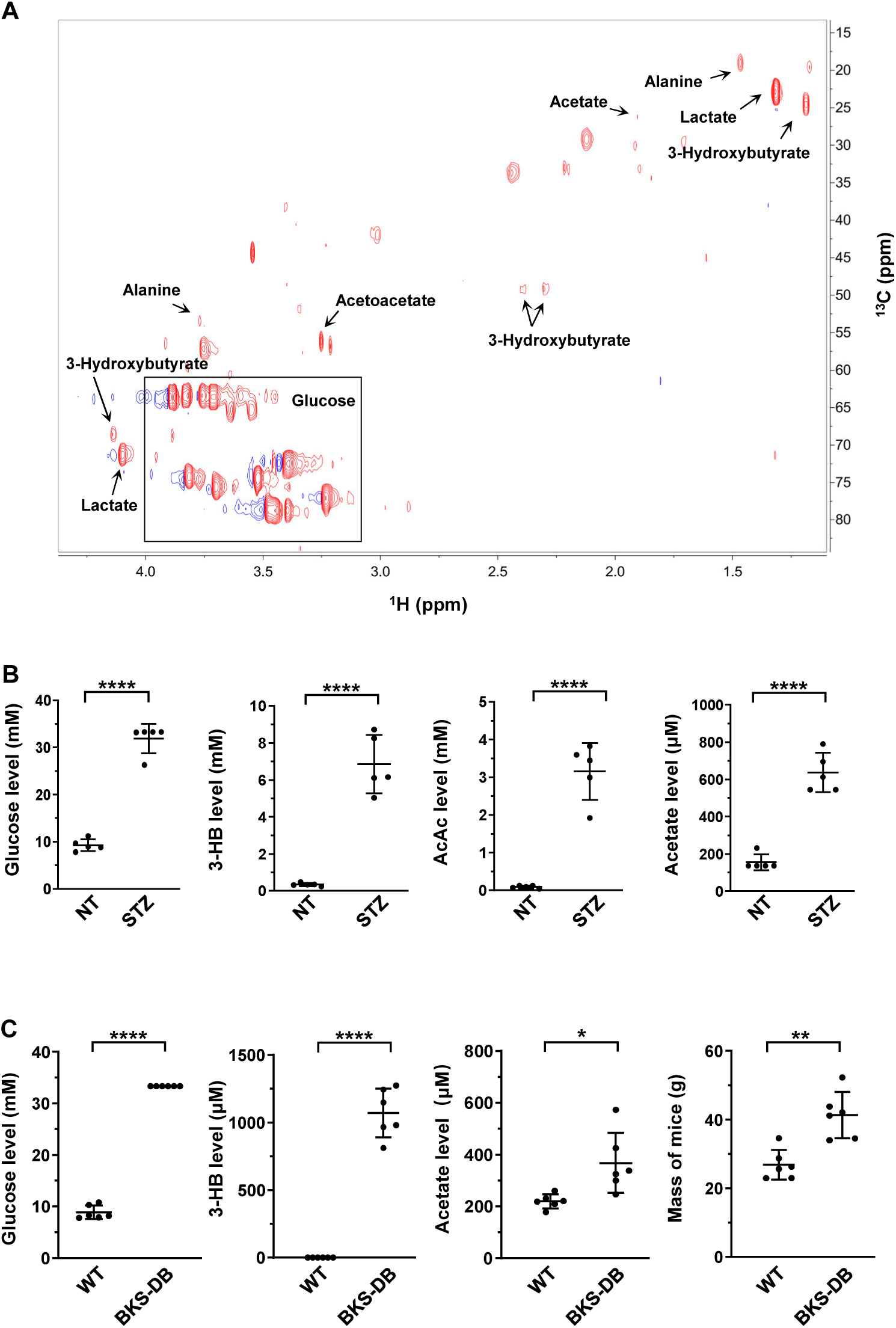
Increased level of acetate in diabetes mellitus. (**A**) Typical 2D ^1^H-^13^C HSQC spectrum of clinical serum sample. (**B**) The levels of serum glucose, 3-HB, AcAc and acetate in STZ-induced diabetic mice (BALB/c, n=5 per group). (**C**) The levels of serum glucose, 3-HB and acetate in db/db mice (C57BL/6, n=6 per group). Abbreviations: 3-HB, 3-hydroxybutyrate; AcAc, acetoacetate; STZ, streptozotocin. Values in (B, C) are expressed as mean±SD and analyzed statistically by two-tailed unpaired Student’s *t* test (\**P*<0.05, \*\**P*<0.01, \*\*\**P*<0.001, \*\*\*\**P*<0.0001, n.s., no significant difference).

**Figure 1—figure supplement 2.**
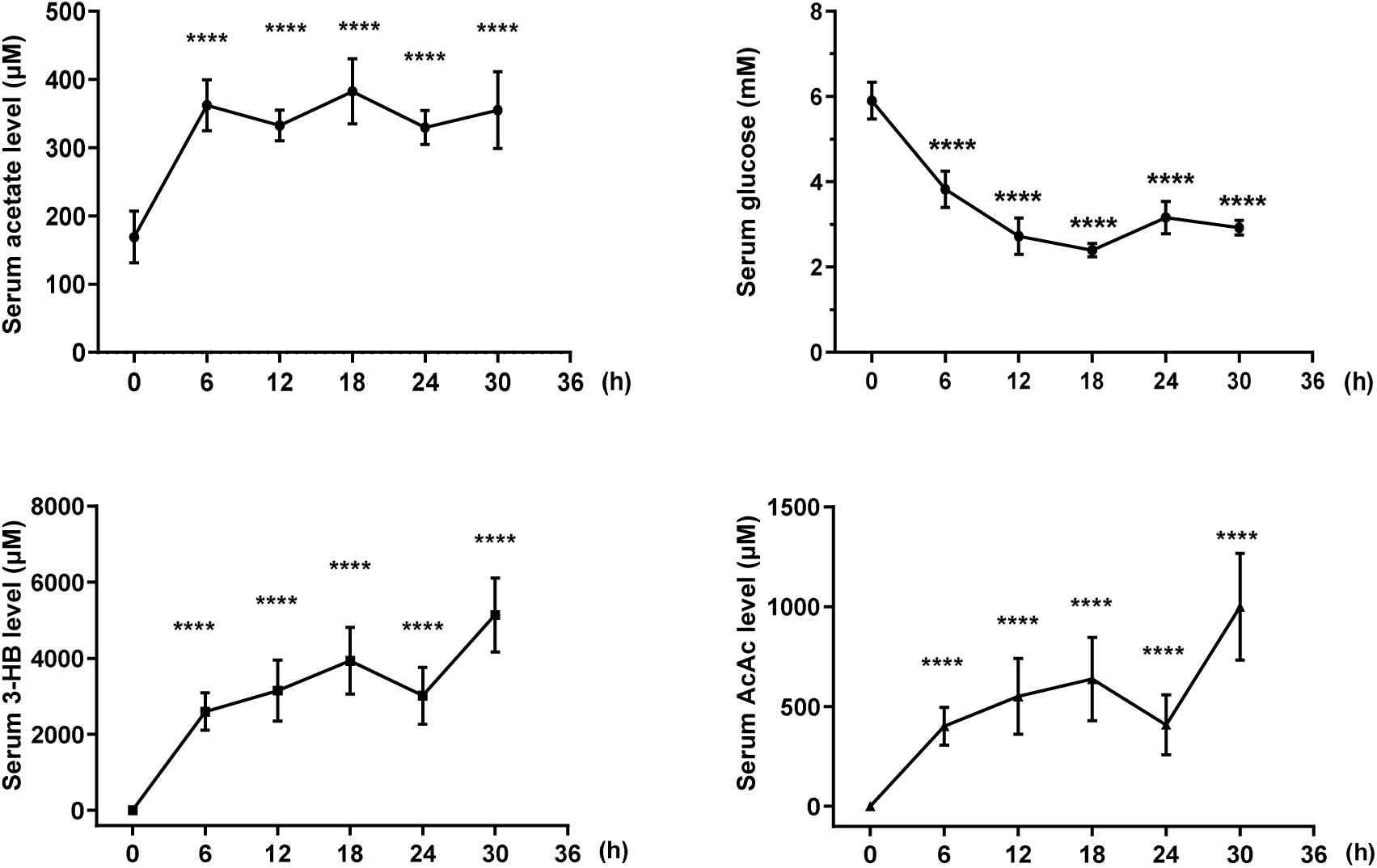
Increased level of acetate in fasting mice. The levels of acetate, 3-HB, AcAc and glucose in the serum of BALB/c mice starved for indicated time course. Values are presented as mean±SD (n=5). Statistics were performed employing two-tailed unpaired Student’s *t* test (\**P*<0.05, \*\**P*<0.01, \*\*\**P*<0.001, \*\*\*\**P*<0.0001, n.s., no significant difference).

**Figure 2—figure supplement 1.**
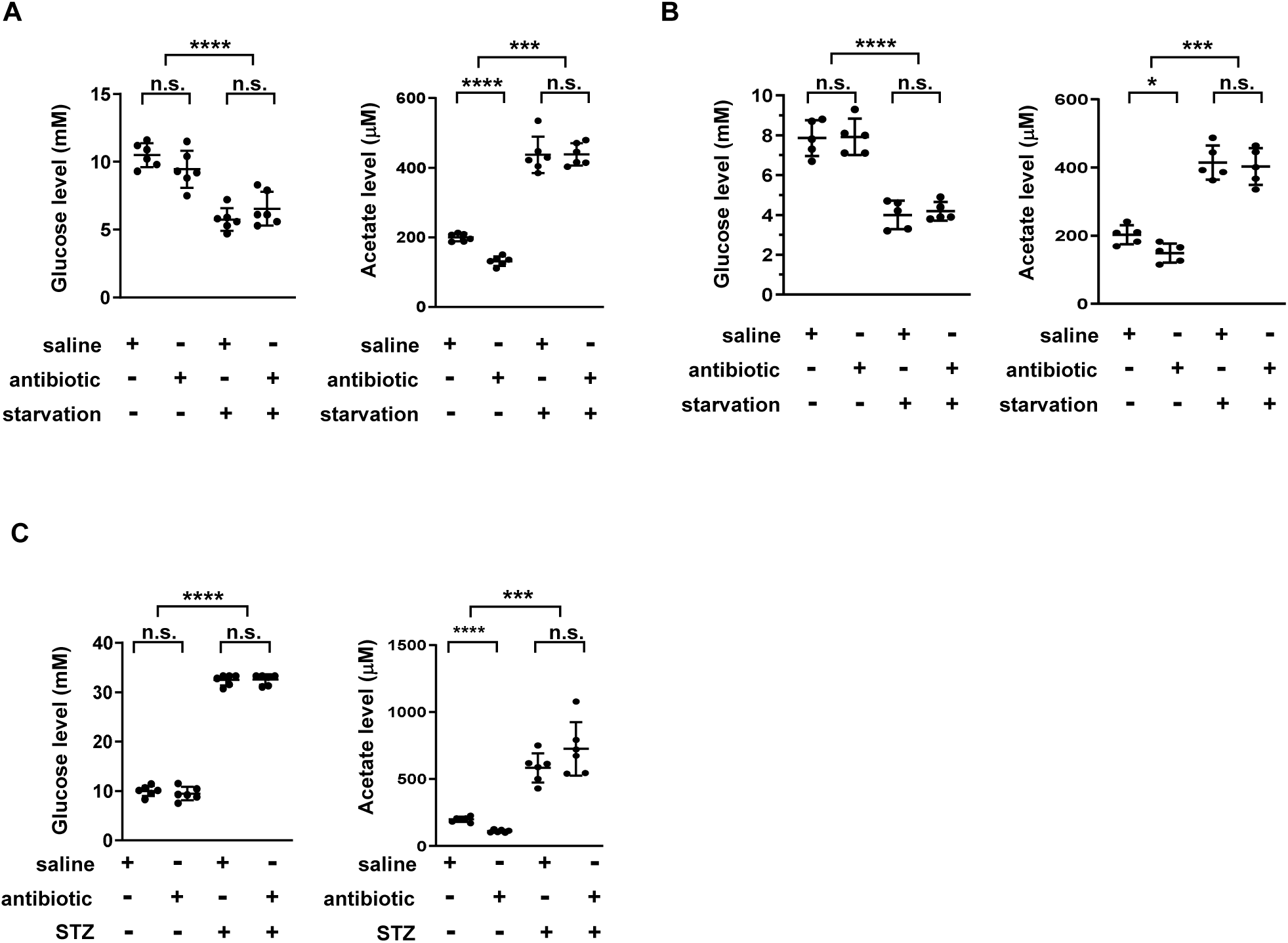
Acetate is increased independently of gut microbiota upon energy stress. (**A**) The levels of serum glucose and acetate of C57BL/6 mice pretreated with antibiotics for 3-weeks and then starved for additional 24 h (n=5). (**B**) The levels of serum glucose and acetate of BALB/c mice pretreated with antibiotics for 3-weeks and then starved for another 12 h (n=5). (**C**) Serum glucose and acetate levels of STZ-induced diabetic mice pretreated with antibiotics for 3-weeks (C57BL/6, n=5). Values are expressed as mean±SD. Statistical analyses were carried out by using two-tailed unpaired Student’s t test (\**P*<0.05, \*\**P*<0.01, \*\*\**P*<0.001, \*\*\*\**P*<0.0001, n.s., no significant difference).

**Figure 2—figure supplement 2.**
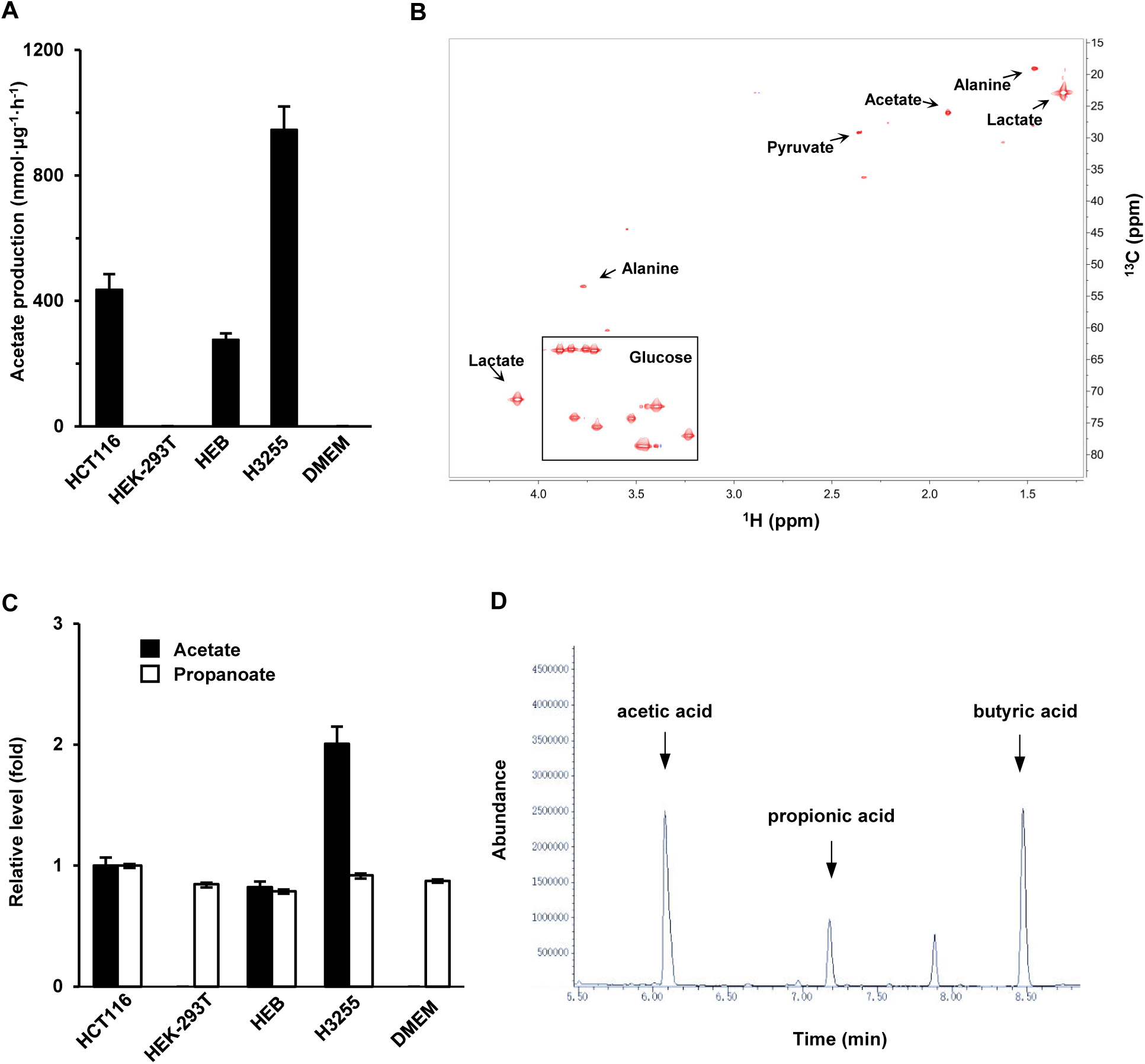
Acetate is secreted by in vitro cultured mammalian cells. (**A**) Acetate secreted by various cells cultured in fresh DMEM medium for 20 h was detected by employing NMR. (**B**) A typical NMR 2D ^1^H-^13^C HSQC spectrum of culture medium in which HCT116 cells were culture for 20 h. (**C**, **D**) The same cells as in (A) were detected for the production of acetate, propanoate and butyrate with GC-MS. Propanoate and butyrate were used as an internal control. Values in (A, C) are expressed as mean±SD (n=3) of three independent experiments.

**Figure 2—figure supplement 3.**
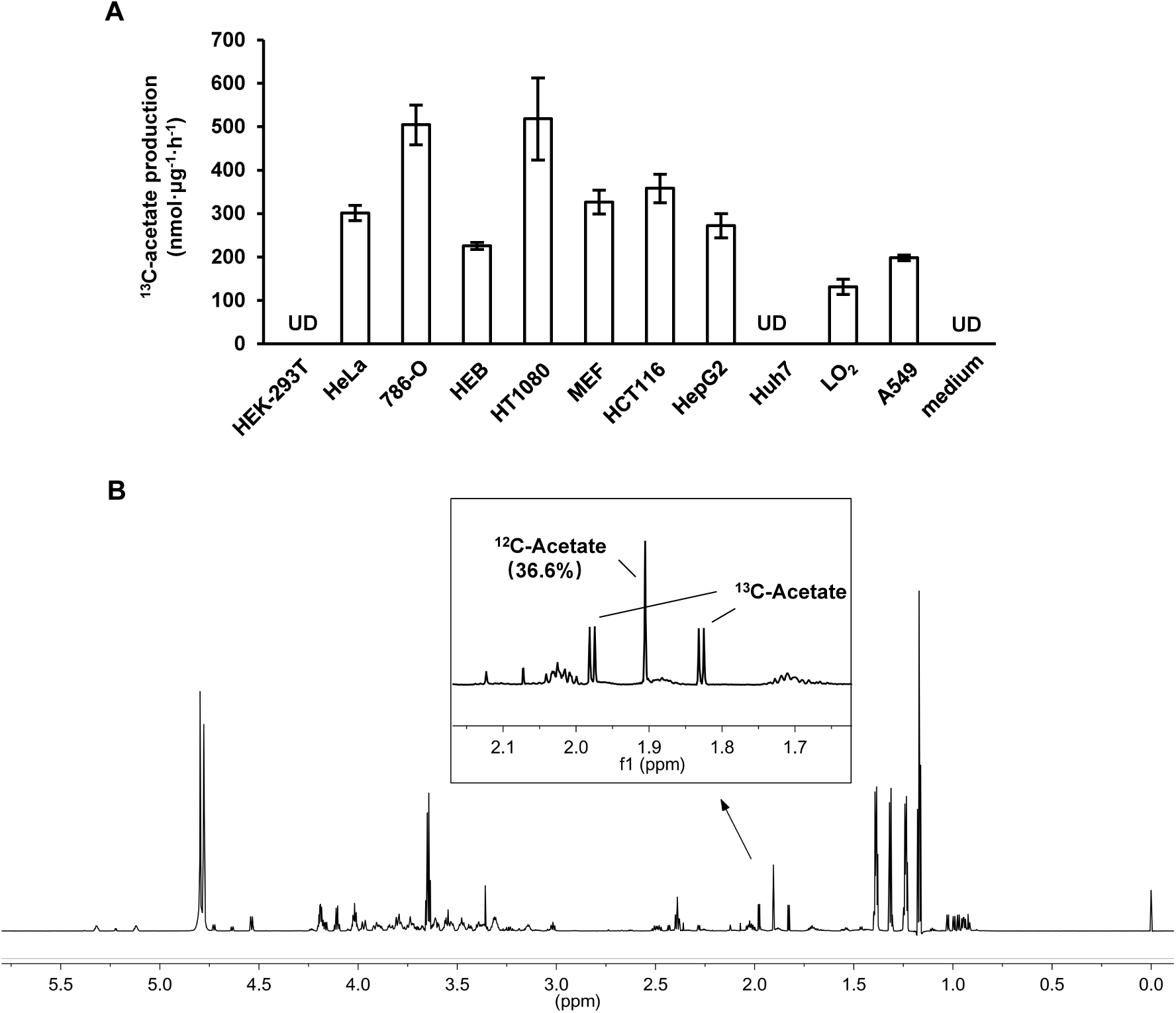
Acetate is derived from other nutrients besides glucose. (**A**) The amount of U-^13^C-acetate secreted by indicated cells cultured in U-^13^C-glucose-containing medium for 20 h. Values are expressed as mean±SD (n=3) of three independent experiments. UD, undetectable. (**B**) 1D ^1^H NMR CPMG spectra of aqueous extracts from medium of LO_2_ cells cultured with U-^13^C-glucose for 20 h.

**Figure 2—figure supplement 4.**
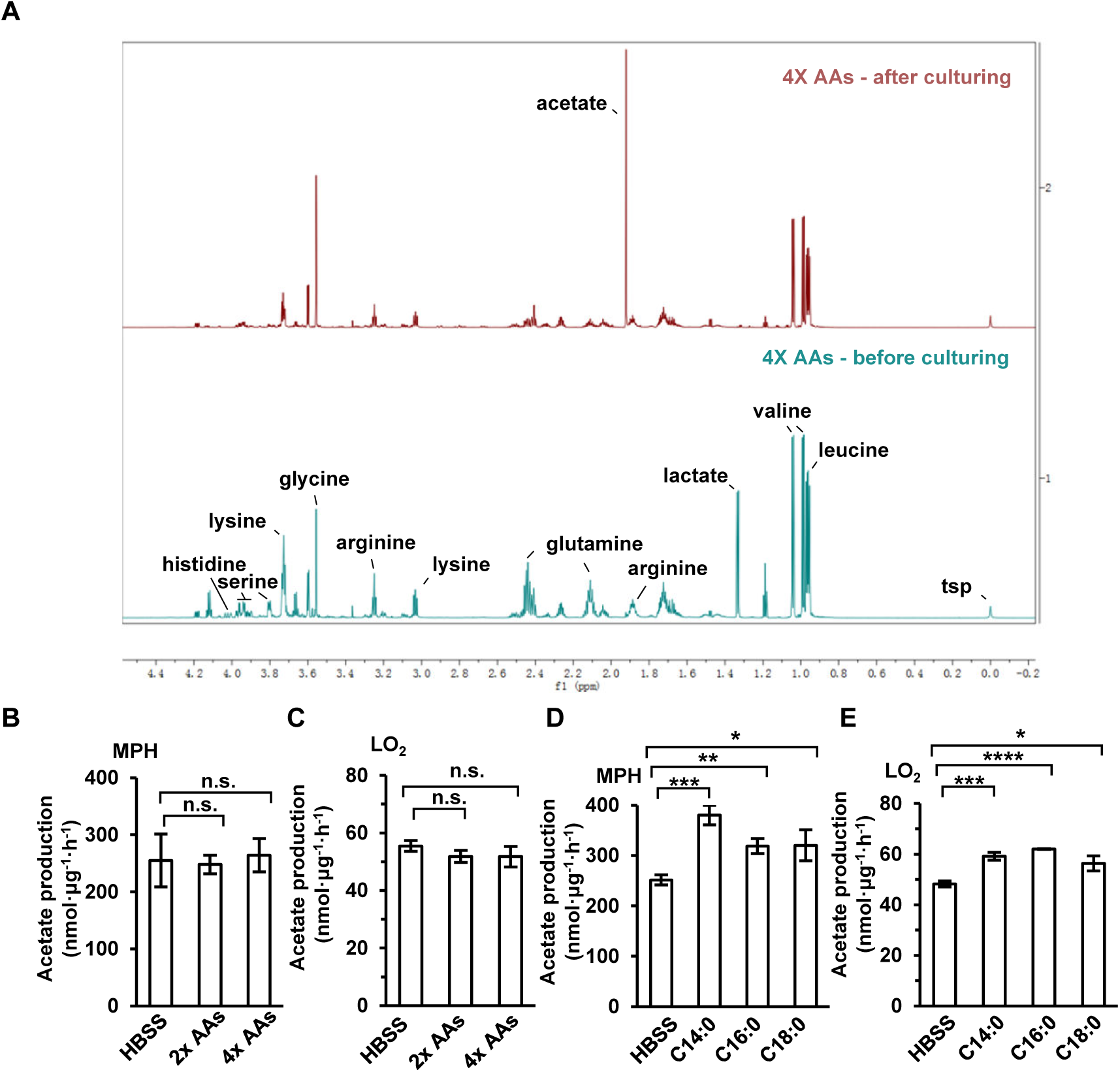
ES-Acetate is mainly derived from FFAs. (**A**) A representative 1D ^1^H NMR CPMG spectra of aqueous extracts from the DMEM medium with 4×AAs in which LO_2_ cells were cultured for 20 h. (**B**, **C**) The acetate production of primary hepatocyte (MPH) (B) and LO_2_ (C) cells cultured in Hanks’ balanced salt solution (HBSS, free for glucose, fatty acids and amino acids) supplemented with or without indicated doses of amino acids (AAs) for 20 h. (**D**, **E**) The acetate production of MPH (D) and LO_2_ (E) cells cultured in HBSS with or without 500 μM of indicated free fatty acids (FFAs) (C14:0, Myristate; C16:0, Palmitate; C18:0, Stearate) for 20 h. Values are expressed as mean±SD (n=3) of three independent measurements for (B-E). Statistics in (B-E) were analyzed by using two-tailed unpaired Student’s *t* test (\**P*<0.05, \*\**P*<0.01, \*\*\**P*<0.001, \*\*\*\**P*<0.0001, n.s., no significant difference).

**Figure 3—source data 1**

Complete, unedited immunoblots, as well as immunoblots including sample and band identification, are provided for the immunoblots presented in Figure 3.

**Figure 3—figure supplement 1.**
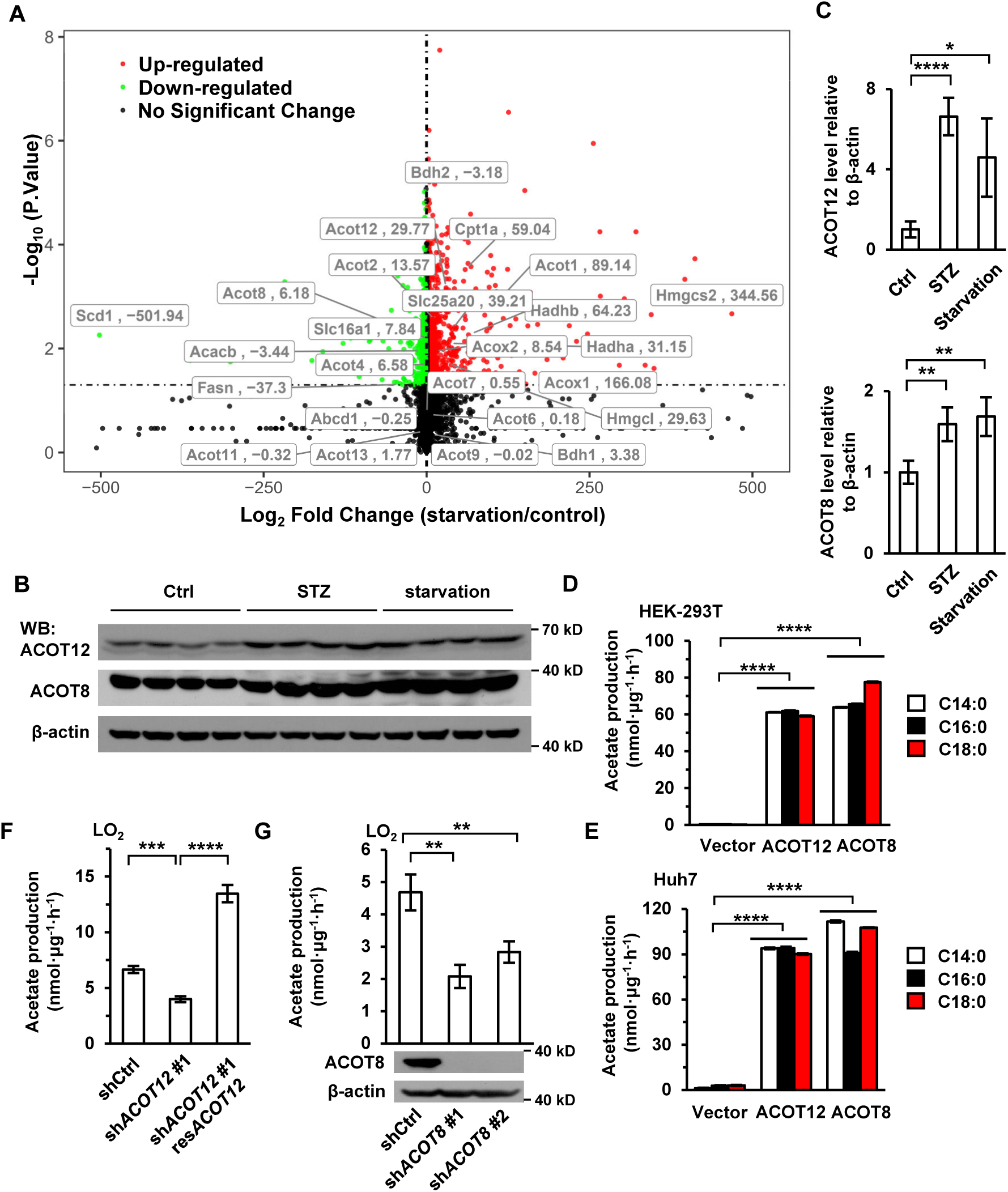
ACOT12 and ACOT8 are responsible for acetate production during energy stress. (**A**) Volcano plot of RNAseq analysis data from Goldstein et al. (2017) (Goldstein et al., 2017). Taxa with fold change >2 and p-value< 0.05 are labeled in red and taxa with fold change < -2 and p-value < 0.05 are labeled in green. (**B**, **C**) ACOT12 and ACOT8 in livers of C57BL/6 mice with STZ-induced type I diabetes or 48 h starvation were detected by Western Blot (B) and their protein levels relative to β-actin are analyzed (C). ACOT, acyl-CoA thioesterase. (**D**, **E**) HEK-293T (D) and Huh7 (E) cell lines overexpressing ACOT12 or ACOT8 were cultured in medium containing indicated FFAs for 20 h, followed by detection of acetate secretion (n=3). (**F**) The enrichment of U- ^13^C-acetate in LO_2_ cells knocked down and further rescued for ACOT12 expression and then cultured in medium supplemented with U-^13^C- palmitate for 20 h (n=3). (**G**) The enrichment of U-^13^C-acetate in LO_2_ cell lines knocked down for ACOT8 and cultured in medium containing U-^13^C-palmitate for 20 h (n=3). Values are expressed as mean±SD of three independent experiments. \**P*<0.05, \*\**P*<0.01, \*\*\**P*<0.001, \*\*\*\**P*<0.0001 by two-tailed unpaired Student’s *t* test.

**Figure 3—figure supplement 1—source data 1**

Complete, unedited immunoblots, as well as immunoblots including sample and band identification, are provided for the immunoblots presented in Figure 3—figure supplement 1.

**Figure 4—source data 1**

Complete, unedited immunoblots, as well as immunoblots including sample and band identification, are provided for the immunoblots presented in Figure 4.

**Figure 4—figure supplement 1.**
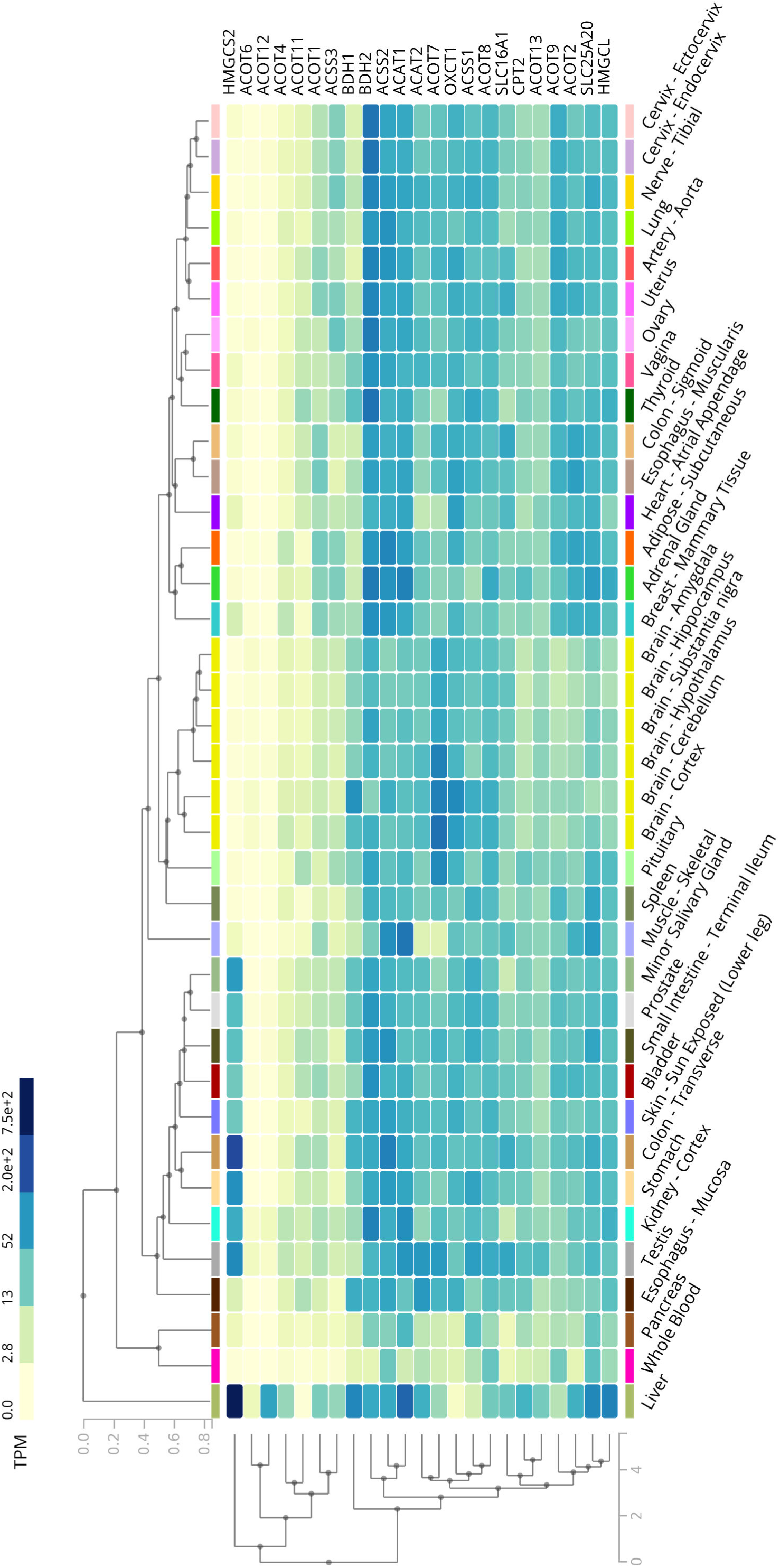
The expression profile of ACOTs and ketogenetic enzymes in human liver. Heatmap from GTEx database represents the expression levels of ACOTs and ketogenetic genes in a variety of normal human tissues.

**Figure 4—figure supplement 2.**
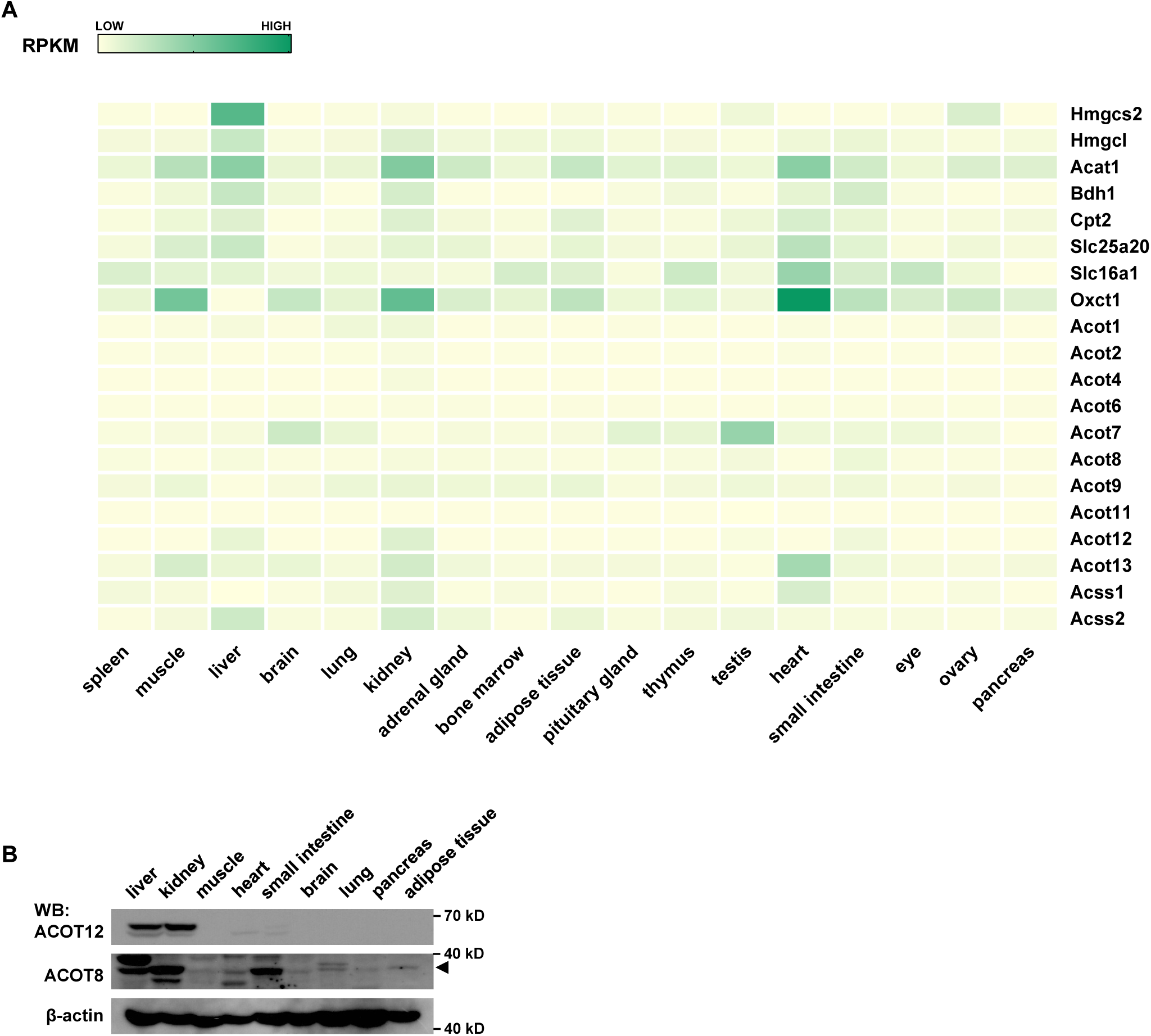
The expression profile of ACOTs and ketogenetic enzymes in mouse liver. **(A)** Heatmap from GEO database represents the expression levels of ACOTs and ketogenetic genes in normal mouse tissues indicated. (**B**) The protein levels of ACOT12 and ACOT8 in different tissues of mice (C57BL/6).

**Figure 4—figure supplement 2—source data 1**

Complete, unedited immunoblots, as well as immunoblots including sample and band identification, are provided for the immunoblots presented in Figure 4—figure supplement 2.

**Figure 4—figure supplement 3.**
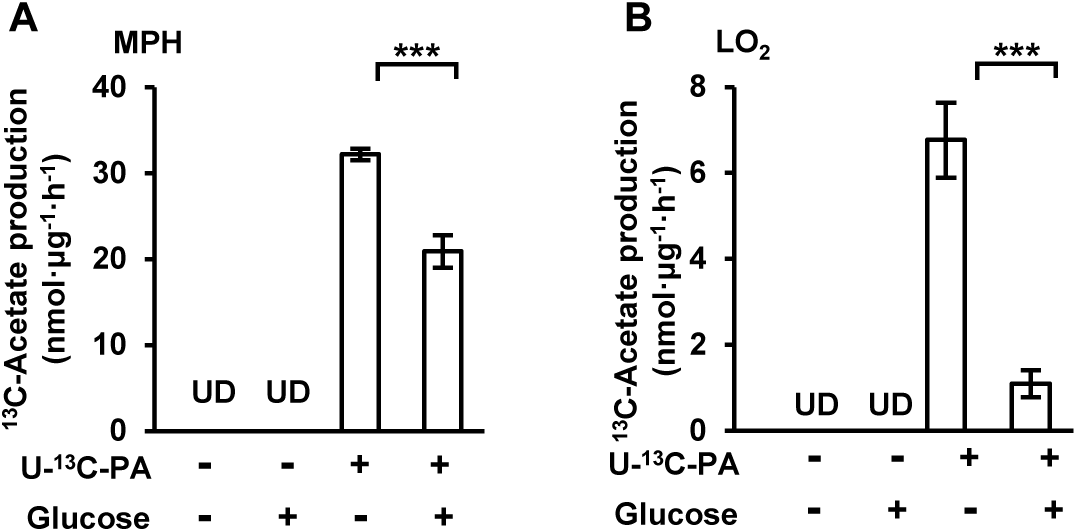
FFA-derived acetate is diminished by supplementation of glucose. (**A**, **B**) NMR detection of the amount of U-^13^C-acetate secreted by MPH (A) and LO_2_ (B) cell lines after incubation in U-^13^C-palmitate-containing HBSS supplemented with or without (w/wo) glucose (20 mM) for 20 h. UD, undetectable. Values are expressed as mean±SD (n=3) of three independent measurements. \**P*<0.05, \*\**P*<0.01, \*\*\**P*<0.001, \*\*\*\**P*<0.0001 by two-tailed unpaired Student’s *t* test.

**Figure 5—source data 1**

Complete, unprocessed immunoblots displaying sample and band identification are presented in Figure 5, along with the corresponding raw data for immunostaining.

**Figure 6—figure supplement 1.**
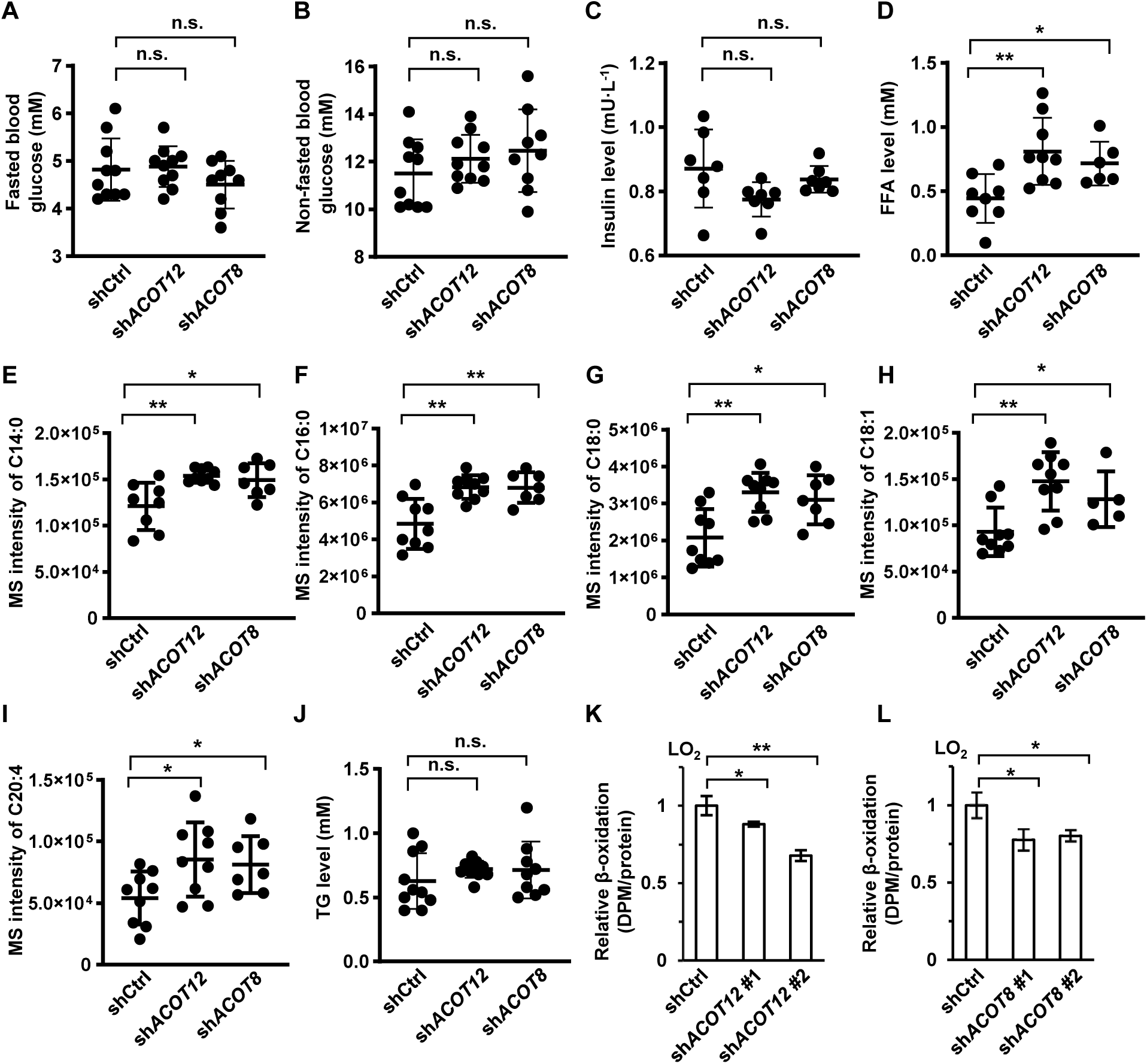
ACOT12 and ACOT8 are involved in the catabolism of fatty acid. (**A**-**D**) Serum levels of fasted blood glucose (A), non-fasted blood glucose (B), insulin and total FFA (D) of C57BL/6 mice with adenovirus-mediated knockdown of ACOT12 or ACOT8. Fasted, mice were fasted for 12 h; Non-fasted, mice were fed normally. (**E**-**I**) Serum free fatty acids’ levels determined by GC-MS. (**J**) Serum levels of triacylglycerol (TG) of C57BL/6 mice with adenovirus-mediated knockdown of ACOT12 or ACOT8. (**K**, **L**) LO_2_ cell lines knocked down for ACOT12 (K) or ACOT8 (L) were cultured in glucose free reaction buffer containing 0.8 μCi/mL [9,10- ^3^H(N)]-oleic acid for 20 h, followed by determination of the relative β- oxidation rate (n=3). Results are expressed as mean±SD of n=10 mice per group in (A-J) and three independent experiments in (K, L), and analyzed by using unpaired Student’s *t* test (\**P*<0.05, \*\**P*<0.01, \*\*\**P*<0.001, \*\*\*\**P*<0.0001, n.s., no significant difference).

**Figure 6—figure supplement 2.**
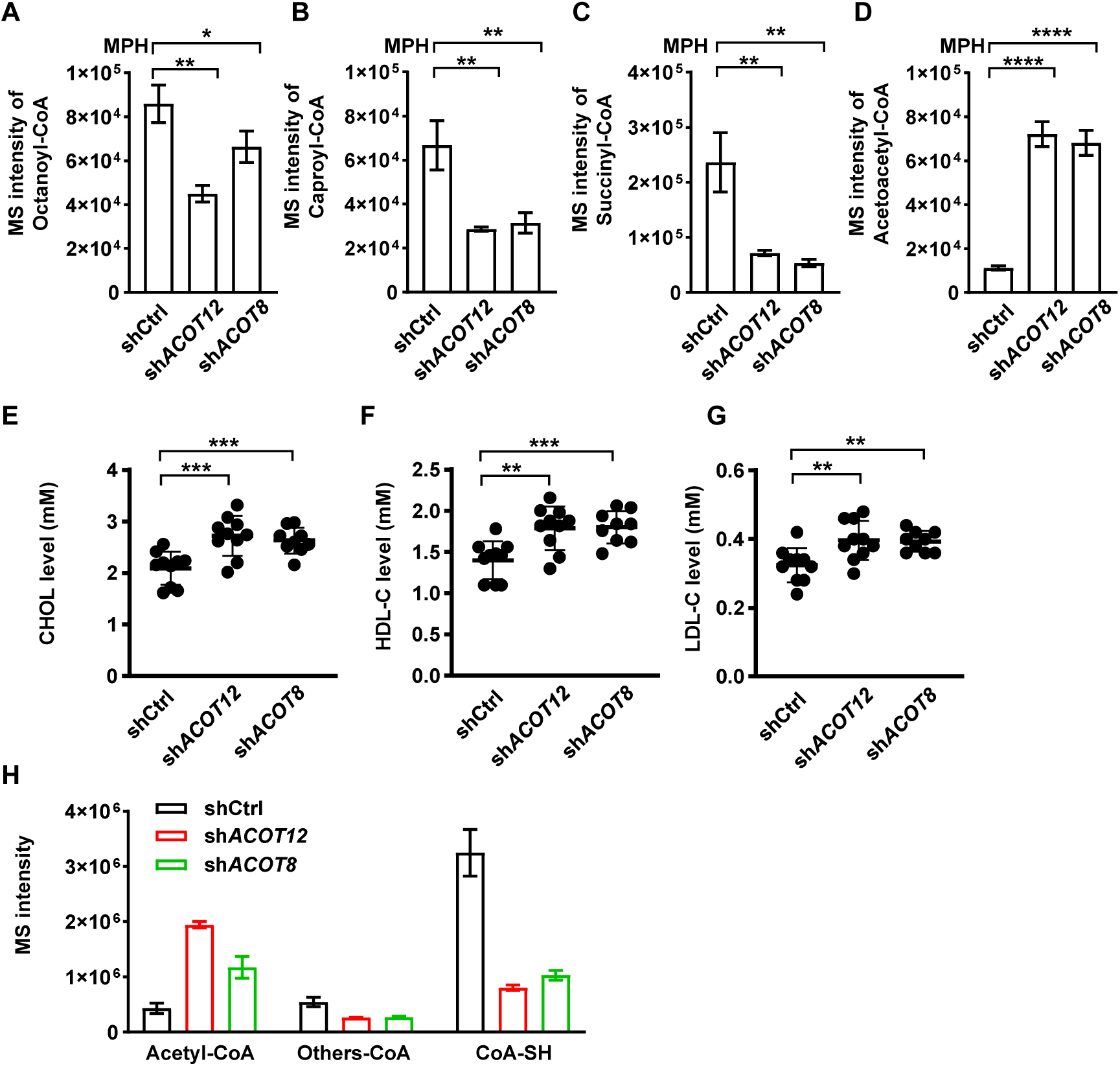
ACOT12 and ACOT8 serve to maintain CoA pool for sustained FAO. (**A**-**D**) Relative abundance of octanoyl-CoA (A), caproyl-CoA (B), succinyl-CoA (C) and acetoacetyl-CoA (D) in MPHs knocked down for ACOT12 or ACOT8 (n=3). (**E**- **G**) Serum levels of cholesterol (CHOL) (E), high density lipoprotein cholesterol (HDL-C) (F) and low density lipoprotein cholesterol (LDL-C) (G) of C57BL/6 mice with adenovirus-mediated knockdown of ACOT12 or ACOT8. (**H**) Relative abundance of reduced CoA, acetyl-CoA and other oxidized CoA (octanoyl-CoA, caproyl-CoA, succinyl-CoA, acetoacetyl-CoA and HMG-CoA) in MPHs knocked down for ACOT12 or ACOT8. HMG-CoA, 3-hydroxy-3-methylglutaryl-CoA (n=3). Results are expressed as mean±SD of three independent experiments in (A-D, H) and n=10 mice per group in (E-G), and analyzed by using unpaired Student’s *t* test (\**P*<0.05, \*\**P*<0.01, \*\*\**P*<0.001, \*\*\*\**P*<0.0001, n.s., no significant difference).

**Figure 7—source data 1**

Complete, unedited immunoblots, as well as immunoblots including sample and band identification, are provided for the immunoblots presented in Figure 7.

**Figure 8—figure supplement 1.**
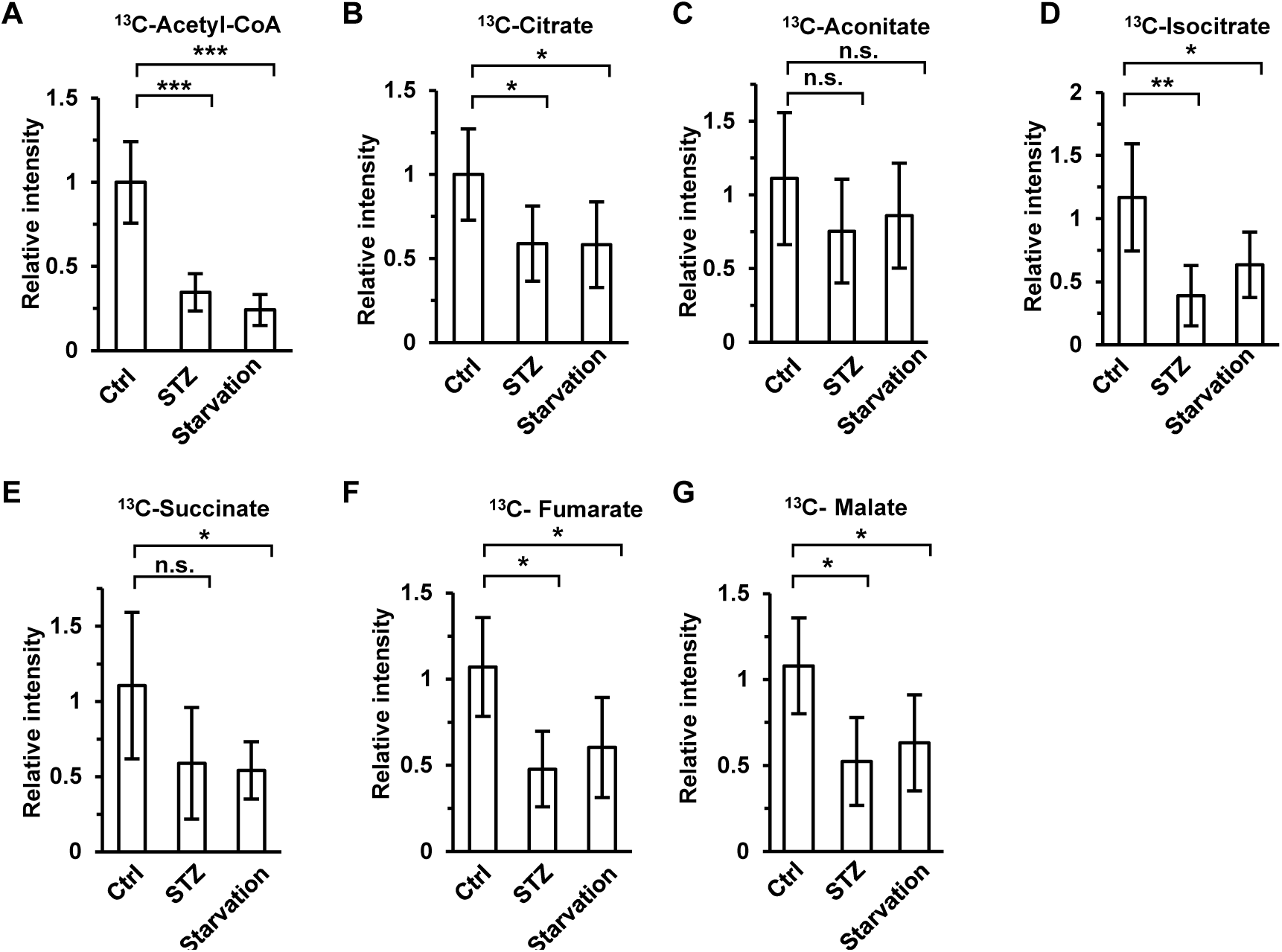
Accumulation of acetate derivatives in muscle is retarded under energy stress. Relative abundance of ^13^C-acetyl-CoA (A), ^13^C-citrate (B), ^13^C-aconitate (C), ^13^C- isocitrate (D), ^13^C-succinate (E), ^13^C-fumarate (F) and ^13^C-malate (G) in the muscle of diabetic and starved mice (C57BL/6) was determined 1 h after intraperitoneal injection of 2-^13^C-acetate (310 mg/kg). Values are expressed as mean±SD of (n=5 mice per group). \**P*<0.05, \*\**P*<0.01, \*\*\**P*<0.001, \*\*\*\**P*<0.0001 and n.s. *P*≥ 0.05 by unpaired Student’s *t* test.

**Figure 8—figure supplement 2.**
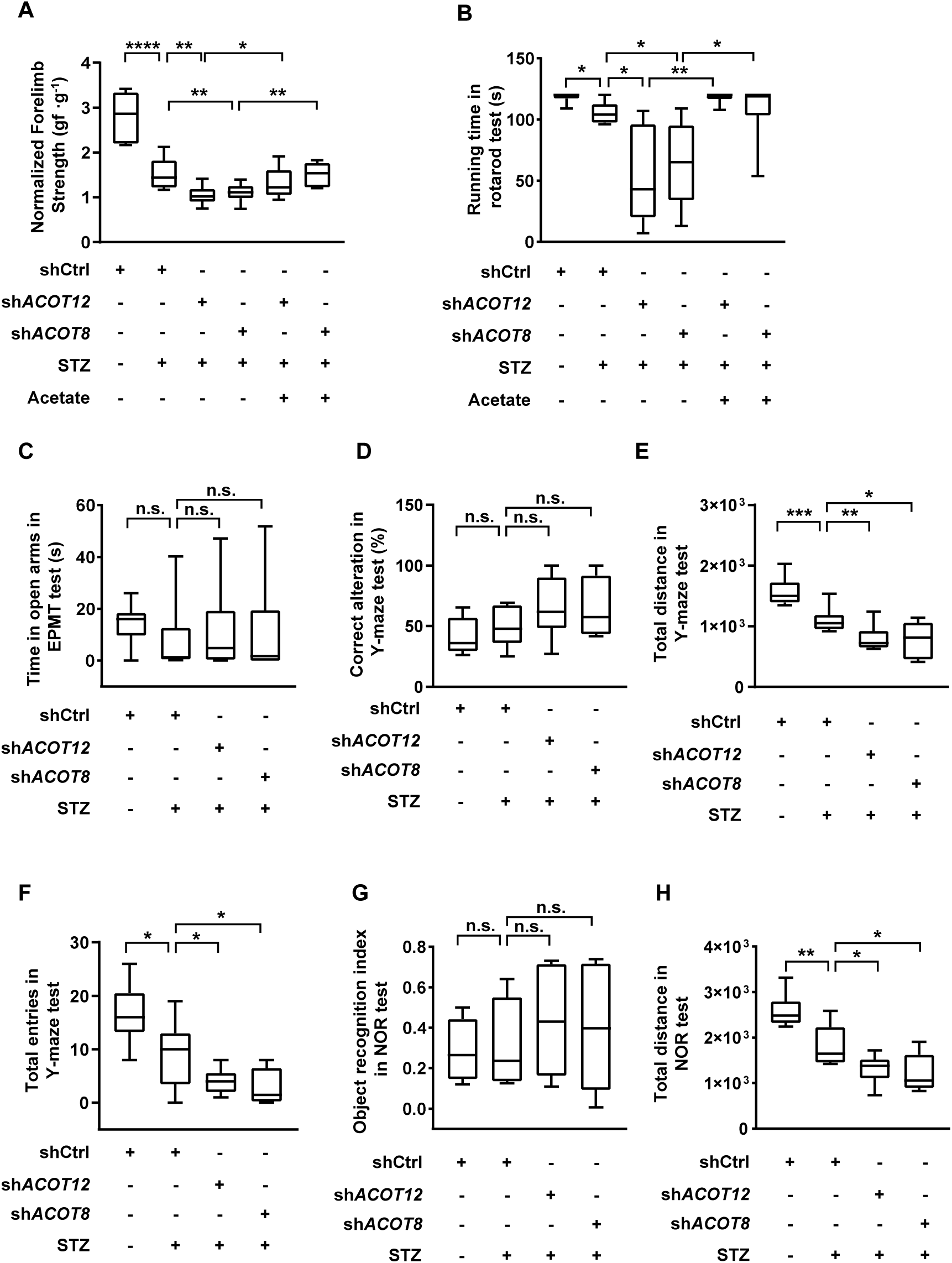
Behavior analyses of diabetic mice with KD of ACOT12 or ACOT8. (**A**, **B**) Normalized forelimb strength in forelimb grip force test (A) and total running time in the rotarod test (B) were determined in STZ-induced diabetic C57BL/6 mice which were knocked down for ACOT12 or ACOT8 in liver and injected intraperitoneally w/wo acetate (300 mg/kg). (**C**) Total time spent in the open arms during the elevated plus maze test of the diabetic C57BL/6 mice w/wo adenovirus- mediated knockdown of ACOT12 or ACOT8 in liver. (**D**-**F**) The percentage of correct alterations (D), total distance moved (E) and total number of entries into each arm (F) in the Y-maze test in the same mice as in (C). (**G**, **H**) The novel object preference index (G) and total distance travelled (H) in NOR test were determined in the same mice as in (C). Results in (A-H) are expressed as box-plot (box extending from the 25th to 75th percentiles with whiskers indicating the minimum and maximum, and lines in boxes indicating the median) of n=10 mice per group, and analyzed by using unpaired Student’s *t* test (\**P*<0.05, \*\**P*<0.01, \*\*\**P*<0.001, \*\*\*\**P*<0.0001, n.s., no significant difference).

## Notes

### Competing Interest Statement

The authors have declared no competing interest.

### Summary of Updates

Section on Materials and methods updated to clarify additional information about clinical trials and animal studies; Figure 1 revised; Figure 6 figure supplement 1 revised; Based on the suggestions from the reviewers, we have made revisions to rectify certain misleading statements in the manuscript.

## References

Akanji, A.O., Humphreys, S., Thursfield, V., and Hockaday, T.D. (1989). The relationship of plasma acetate with glucose and other blood intermediary metabolites in non-diabetic and diabetic subjects. Clin Chim Acta 185, 25–34.

Cahill, G.F. (2006). Fuel metabolism in starvation. Annu Rev Nutr 26, 1–22.

Castro, M.A., Beltran, F.A., Brauchi, S., and Concha, II (2009). A metabolic switch in brain: glucose and lactate metabolism modulation by ascorbic acid. J Neurochem 110, 423–440.

Cerdan, S., Rodrigues, T.B., Sierra, A., Benito, M., Fonseca, L.L., Fonseca, C.P., and Garcia-Martin, M.L. (2006). The redox switch/redox coupling hypothesis. Neurochem Int 48, 523–530.

Comerford, S.A., Huang, Z., Du, X., Wang, Y., Cai, L., Witkiewicz, A.K., Walters, H., Tantawy, M.N., Fu, A., Manning, H.C., et al. (2014). Acetate dependence of tumors. Cell 159, 1591–1602.

D’Acunzo, P., Perez-Gonzalez, R., Kim, Y., Hargash, T., Miller, C., Alldred, M.J., Erdjument-Bromage, H., Penikalapati, S.C., Pawlik, M., Saito, M., et al. (2021). Mitovesicles are a novel population of extracellular vesicles of mitochondrial origin altered in Down syndrome. Sci Adv 7.

Denechaud, P.D., Girard, J., and Postic, C. (2008). Carbohydrate responsive element binding protein and lipid homeostasis. Curr Opin Lipidol 19, 301–306.

Dentin, R., Denechaud, P.D., Benhamed, F., Girard, J., and Postic, C. (2006). Hepatic gene regulation by glucose and polyunsaturated fatty acids: a role for ChREBP. J Nutr 136, 1145–1149.

Fagerberg, L., Hallstrom, B.M., Oksvold, P., Kampf, C., Djureinovic, D., Odeberg, J., Habuka, M., Tahmasebpoor, S., Danielsson, A., Edlund, K., et al. (2014). Analysis of the human tissue-specific expression by genome-wide integration of transcriptomics and antibody-based proteomics. Mol Cell Proteomics 13, 397–406.

Fellows, R., Denizot, J., Stellato, C., Cuomo, A., Jain, P., Stoyanova, E., Balazsi, S., Hajnady, Z., Liebert, A., Kazakevych, J., et al. (2018). Microbiota derived short chain fatty acids promote histone crotonylation in the colon through histone deacetylases. Nat Commun 9, 105.

Field, A.E., Coakley, E.H., Must, A., Spadano, J.L., Laird, N., Dietz, W.H., Rimm, E., and Colditz, G.A. (2001). Impact of overweight on the risk of developing common chronic diseases during a 10-year period. Arch Intern Med 161, 1581–1586.

Fruhbeck, G., Gomez-Ambrosi, J., Muruzabal, F.J., and Burrell, M.A. (2001). The adipocyte: a model for integration of endocrine and metabolic signaling in energy metabolism regulation. Am J Physiol-Endoc M 280, E827–E847.

Galgani, J., and Ravussin, E. (2008). Energy metabolism, fuel selection and body weight regulation. Int J Obesity 32, S109–S119.

Goldberg, I.J., Reue, K., Abumrad, N.A., Bickel, P.E., Cohen, S., Fisher, E.A., Galis, Z.S., Granneman, J.G., Lewandowski, E.D., Murphy, R., et al. (2018). Deciphering the Role of Lipid Droplets in Cardiovascular Disease: A Report From the 2017 National Heart, Lung, and Blood Institute Workshop. Circulation 138, 305–315.

Goldstein, I., Baek, S., Presman, D.M., Paakinaho, V., Swinstead, E.E., and Hager, G.L. (2017). Transcription factor assisted loading and enhancer dynamics dictate the hepatic fasting response. Genome Res 27, 427–439.

Gonzalez, C., Cuvellier, S., Hue-Beauvais, C., and Levi-Strauss, M. (2003). Genetic control of non obese diabetic mice susceptibility to high-dose streptozotocin-induced diabetes. Diabetologia 46, 1291–1295.

He, T.C., Zhou, S.B., da Costa, L.T., Yu, J., Kinzler, K.W., and Vogelstein, B. (1998). A simplified system for generating recombinant adenoviruses. P Natl Acad Sci USA 95, 2509–2514.

Hirai, S., Miwa, H., Tanaka, T., Toriumi, K., Kunii, Y., Shimbo, H., Sakamoto, T., Hino, M., Izumi, R., Nagaoka, A., et al. (2021). High-sucrose diets contribute to brain angiopathy with impaired glucose uptake and psychosis-related higher brain dysfunctions in mice. Sci Adv 7, eabl6077.

Huang, P.Y., He, Z.Y., Ji, S.Y., Sun, H.W., Xiang, D., Liu, C.C., Hu, Y.P., Wang, X., and Hui, L.J. (2011). Induction of functional hepatocyte-like cells from mouse fibroblasts by defined factors. Nature 475, 386–U142.

Hui, S., Cowan, A.J., Zeng, X., Yang, L., TeSlaa, T., Li, X., Bartman, C., Zhang, Z., Jang, C., Wang, L., et al. (2020). Quantitative Fluxomics of Circulating Metabolites. Cell Metab 32, 676–688 e674.

Ishizuka, M., Toyama, Y., Watanabe, H., Fujiki, Y., Takeuchi, A., Yamasaki, S., Yuasa, S., Miyazaki, M., Nakajima, N., Taki, S., et al. (2004). Overexpression of human acyl-CoA thioesterase upregulates peroxisome biogenesis. Exp Cell Res 297, 127–141.

Krishnakumar, A.M., Sliwa, D., Endrizzi, J.A., Boyd, E.S., Ensign, S.A., and Peters, J.W. (2008). Getting a handle on the role of coenzyme M in alkene metabolism. Microbiol Mol Biol Rev 72, 445–456.

Lazarow, P.B. (1978). Rat liver peroxisomes catalyze the beta oxidation of fatty acids. J Biol Chem 253, 1522–1528.

Leighton, F., Bergseth, S., Rortveit, T., Christiansen, E.N., and Bremer, J. (1989). Free acetate production by rat hepatocytes during peroxisomal fatty acid and dicarboxylic acid oxidation. J Biol Chem 264, 10347–10350.

Li, T.Y., Song, L.T., Sun, Y., Li, J.Y., Yi, C., Lam, S.M., Xu, D.J., Zhou, L.K., Li, X.T., Yang, Y., et al. (2018). Tip60-mediated lipin 1 acetylation and ER translocation determine triacylglycerol synthesis rate. Nature Communications 9.

Like, A.A., and Rossini, A.A. (1976). Streptozotocin-induced pancreatic insulitis: new model of diabetes mellitus. Science 193, 415–417.

Lindsay, D.B., and Setchell, B.P. (1976). The oxidation of glucose, ketone bodies and acetate by the brain of normal and ketonaemic sheep. J Physiol 259, 801–823.

Liu, Q., Zhang, F.G., Zhang, W.S., Pan, A., Yang, Y.L., Liu, J.F., Li, P., Liu, B.L., and Qi, L.W. (2017). Ginsenoside Rg1 Inhibits Glucagon-Induced Hepatic Gluconeogenesis through Akt-FoxO1 Interaction. Theranostics 7, 4001–4012.

Liu, X., Cooper, D.E., Cluntun, A.A., Warmoes, M.O., Zhao, S., Reid, M.A., Liu, J., Lund, P.J., Lopes, M., Garcia, B.A., et al. (2018). Acetate Production from Glucose and Coupling to Mitochondrial Metabolism in Mammals. Cell 175, 502–513 e513.

Lodhi, I.J., and Semenkovich, C.F. (2014). Peroxisomes: a nexus for lipid metabolism and cellular signaling. Cell Metab 19, 380–392.

Lu, M., Zhu, W.W., Wang, X., Tang, J.J., Zhang, K.L., Yu, G.Y., Shao, W.Q., Lin, Z.F., Wang, S.H., Lu, L., et al. (2019). ACOT12-Dependent Alteration of Acetyl-CoA Drives Hepatocellular Carcinoma Metastasis by Epigenetic Induction of Epithelial-Mesenchymal Transition. Cell Metab 29, 886–900 e885.

Martinic, M.M., and von Herrath, M.G. (2008). Real-time imaging of the pancreas during development of diabetes. Immunol Rev 221, 200–213.

Mashimo, T., Pichumani, K., Vemireddy, V., Hatanpaa, K.J., Singh, D.K., Sirasanagandla, S., Nannepaga, S., Piccirillo, S.G., Kovacs, Z., Foong, C., et al. (2014). Acetate is a bioenergetic substrate for human glioblastoma and brain metastases. Cell 159, 1603–1614.

Meier, U., and Gressner, A.M. (2004). Endocrine regulation of energy metabolism: Review of pathobiochemical and clinical chemical aspects of leptin, ghrelin, adiponectin, and resistin. Clin Chem 50, 1511–1525.

Must, A., Spadano, J., Coakley, E.H., Field, A.E., Colditz, G., and Dietz, W.H. (1999). The disease burden associated with overweight and obesity. Jama-J Am Med Assoc 282, 1523–1529.

Nishimoto, S., Fukuda, D., Higashikuni, Y., Tanaka, K., Hirata, Y., Murata, C., Kim-Kaneyama, J.R., Sato, F., Bando, M., Yagi, S., et al. (2016). Obesity-induced DNA released from adipocytes stimulates chronic adipose tissue inflammation and insulin resistance. Sci Adv 2, e1501332.

Palikaras, K., Lionaki, E., and Tavernarakis, N. (2015). Balancing mitochondrial biogenesis and mitophagy to maintain energy metabolism homeostasis. Cell Death Differ 22, 1399–1401.

Puchalska, P., and Crawford, P.A. (2017). Multi-dimensional Roles of Ketone Bodies in Fuel Metabolism, Signaling, and Therapeutics. Cell Metabolism 25, 262–284.

Ritchie, M.E., Phipson, B., Wu, D., Hu, Y., Law, C.W., Shi, W., and Smyth, G.K. (2015). limma powers differential expression analyses for RNA-sequencing and microarray studies. Nucleic Acids Res 43, e47.

Robinson, A.M., and Williamson, D.H. (1980). Physiological Roles of Ketone-Bodies as Substrates and Signals in Mammalian-Tissues. Physiol Rev 60, 143–187.

Russell, J.B., and Cook, G.M. (1995). Energetics of Bacterial-Growth - Balance of Anabolic and Catabolic Reactions. Microbiol Rev 59, 48–62.

Sakakibara, I., Fujino, T., Ishii, M., Tanaka, T., Shimosawa, T., Miura, S., Zhang, W., Tokutake, Y., Yamamoto, J., Awano, M., et al. (2009). Fasting-induced hypothermia and reduced energy production in mice lacking acetyl-CoA synthetase 2. Cell Metab 9, 191–202.

Schug, Z.T., Peck, B., Jones, D.T., Zhang, Q., Grosskurth, S., Alam, I.S., Goodwin, L.M., Smethurst, E., Mason, S., Blyth, K., et al. (2015). Acetyl-CoA synthetase 2 promotes acetate utilization and maintains cancer cell growth under metabolic stress. Cancer Cell 27, 57–71.

Schug, Z.T., Vande Voorde, J., and Gottlieb, E. (2016). The metabolic fate of acetate in cancer. Nat Rev Cancer 16, 708–717.

Seufert, C.D., Mewes, W., and Soeling, H.D. (1984). Effect of long-term starvation on acetate and ketone body metabolism in obese patients. Eur J Clin Invest 14, 163–170.

Sivan, A., Corrales, L., Hubert, N., Williams, J.B., Aquino-Michaels, K., Earley, Z.M., Benyamin, F.W., Lei, Y.M., Jabri, B., Alegre, M.L., et al. (2015). Commensal Bifidobacterium promotes antitumor immunity and facilitates anti-PD-L1 efficacy. Science 350, 1084–1089.

Sivanand, S., Viney, I., and Wellen, K.E. (2018). Spatiotemporal Control of Acetyl-CoA Metabolism in Chromatin Regulation. Trends Biochem Sci 43, 61–74.

Swarbrick, C.M.D., Roman, N., Cowieson, N., Patterson, E.I., Nanson, J., Siponen, M.I., Berglund, H., Lehtio, L., and Forwood, J.K. (2014). Structural Basis for Regulation of the Human Acetyl-CoA Thioesterase 12 and Interactions with the Steroidogenic Acute Regulatory Protein-related Lipid Transfer (START) Domain. J Biol Chem 289, 24263–24274.

Thorrez, L., Laudadio, I., Van Deun, K., Quintens, R., Hendrickx, N., Granvik, M., Lemaire, K., Schraenen, A., Van Lommel, L., Lehnert, S., et al. (2011). Tissue-specific disallowance of housekeeping genes: the other face of cell differentiation. Genome Res 21, 95–105.

Tillander, V., Alexson, S.E.H., and Cohen, D.E. (2017). Deactivating Fatty Acids: Acyl-CoA Thioesterase-Mediated Control of Lipid Metabolism. Trends Endocrinol Metab 28, 473–484.

Todesco, T., Zamboni, M., Armellini, F., Bissoli, L., Turcato, E., Piemonte, G., Rao, A.V., Jenkins, D.J., and Bosello, O. (1993). Plasma acetate levels in a group of obese diabetic, obese normoglycemic, and control subjects and their relationships with other blood parameters. Am J Gastroenterol 88, 751–755.

Vetizou, M., Pitt, J.M., Daillere, R., Lepage, P., Waldschmitt, N., Flament, C., Rusakiewicz, S., Routy, B., Roberti, M.P., Duong, C.P.M., et al. (2015). Anticancer immunotherapy by CTLA-4 blockade relies on the gut microbiota. Science 350, 1079-+.

Wang, T., Cao, Y., Zheng, Q., Tu, J., Zhou, W., He, J., Zhong, J., Chen, Y., Wang, J., Cai, R., et al. (2019). SENP1-Sirt3 Signaling Controls Mitochondrial Protein Acetylation and Metabolism. Mol Cell 75, 823–834 e825.

