## Supplementary material for "Hepatic conversion of acetyl-CoA to acetate plays crucial roles in energy stresses": Key Resources Table

| **Key Resources Table** | | | | |
| --- | --- | --- | --- | --- |
| **Reagent type (species) or resource** | **Designation** | **Source or reference** | **Identifiers** | **Additional information** |
| gene (*Homo-sapiens*) | ACOT1 | Core Facility of Biomedical Sciences, Xiamen University | gene ID: 25082 |  |
| gene (*Homo-sapiens*) | ACOT2 | Core Facility of Biomedical Sciences, Xiamen University | gene ID: 15824 |  |
| gene (*Homo-sapiens*) | ACOT4 | Core Facility of Biomedical Sciences, Xiamen University | gene ID: 9637 |  |
| gene (*Homo-sapiens*) | ACOT8 | Core Facility of Biomedical Sciences, Xiamen University | gene ID: 24012 |  |
| gene (*Homo-sapiens*) | ACOT9 | Core Facility of Biomedical Sciences, Xiamen University | gene ID: 17595 |  |
| gene (*Homo-sapiens*) | ACOT11 | Core Facility of Biomedical Sciences, Xiamen University | gene ID: 10617 |  |
| gene (*Homo-sapiens*) | ACOT12 | Core Facility of Biomedical Sciences, Xiamen University | gene ID: 134526 |  |
| cell line (*Homo-sapiens*) | HeLa | our laboratory cells bank |  | Cell line maintained in our laboratory cells bank |
| cell line (*Homo-sapiens*) | HEK-293T | our laboratory cells bank |  | Cell line maintained in our laboratory cells bank |
| cell line (*Homo-sapiens*) | HT1080 | our laboratory cells bank |  | Cell line maintained in our laboratory cells bank |
| cell line (*Homo-sapiens*) | Huh7 | our laboratory cells bank |  | Cell line maintained in our laboratory cells bank |
| cell line (*Homo-sapiens*) | LO_2_ | our laboratory cells bank |  | Cell line maintained in our laboratory cells bank |
| cell line (*Homo-sapiens*) | H3255 | our laboratory cells bank |  | Cell line maintained in our laboratory cells bank |
| cell line (*Homo-sapiens*) | A549 | our laboratory cells bank |  | Cell line maintained in our laboratory cells bank |
| cell line (*Homo-sapiens*) | QBI-293A | our laboratory cells bank |  | Cell line maintained in our laboratory cells bank |
| cell line (*Homo-sapiens*) | HEB | our laboratory cells bank |  | Cell line maintained in our laboratory cells bank |
| cell line (*Homo-sapiens*) | HCT116 | Cell Bank of the Chinese Academy of Sciences (Shanghai) |  | Cell line maintained in Cell Bank of the Chinese Academy of Sciences (Shanghai) |
| cell line (*Homo-sapiens*) | 786-O | Cell Bank of the Chinese Academy of Sciences (Shanghai) |  | Cell line maintained in Cell Bank of the Chinese Academy of Sciences (Shanghai) |
| cell line (*Homo-sapiens*) | HepG2 | Cell Bank of the Chinese Academy of Sciences (Shanghai) |  | Cell line maintained in Cell Bank of the Chinese Academy of Sciences (Shanghai) |
| cell line (*mouse*) | Hepa1-6 | Cell Bank of the Chinese Academy of Sciences (Shanghai) |  | Cell line maintained in Cell Bank of the Chinese Academy of Sciences (Shanghai) |
| cell line (*mouse*) | AML12 | Cell Bank of the Chinese Academy of Sciences (Shanghai) |  | Cell line maintained in Cell Bank of the Chinese Academy of Sciences (Shanghai) |
| transfected construct (*human*) | ACLY shRNA-#1 | This paper |  | Lentiviral construct to transfect and express the shRNA;Targeting sequence: GCAGCAGACCTATGACTATGC |
| transfected construct (*human*) | ACLY shRNA-#2 | This paper |  | Lentiviral construct to transfect and express the shRNA; Targeting sequence: GCATCGCAAACTTCACCAACG |
| transfected construct (*human*) | ACLY shRNA-#3 | This paper |  | Lentiviral construct to transfect and express the shRNA; Targeting sequence: GCACGAAGTCACAATCTTTGT |
| transfected construct (*human*) | ACLY shRNA-#4 | This paper |  | Lentiviral construct to transfect and express the shRNA; Targeting sequence: GCAAGGCATGCTGGACTTTGA |
| transfected construct (*human*) | CPT1A shRNA-#3 | This paper |  | Lentiviral construct to transfect and express the shRNA; Targeting sequence: TACAGTCGGTGAGGCCTCTTATGAA |
| transfected construct (*human*) | CPT1A shRNA-#4 | This paper |  | Lentiviral construct to transfect and express the shRNA; Targeting sequence: GGACCAAGATTACAGTGGTATTTGA |
| transfected construct (*human*) | ABCD1 shRNA-#1 | This paper |  | Lentiviral construct to transfect and express the shRNA; Targeting sequence: GCAGATCAACCTCATCCTTCT |
| transfected construct (*human*) | ACOT12 shRNA-#1 | This paper |  | Lentiviral construct to transfect and express the shRNA; Targeting sequence: GCTAGAGTTGGACAAGTTATA |
| transfected construct (*human*) | ACOT12 shRNA-#2 | This paper |  | Lentiviral construct to transfect and express the shRNA; Targeting sequence: CAAATACCAGTGATTTGGATTAGCA |
| transfected construct (*mouse*) | ACOT12 shRNA-#5 | This paper |  | Lentiviral construct to transfect and express the shRNA; Targeting sequence: GCATGGAGATCAGTATCAAGG |
| transfected construct (*mouse*) | ACOT12 shRNA-#6 | This paper |  | Lentiviral construct to transfect and express the shRNA; Targeting sequence: GCAGGTTCAGCGATTCCATTT |
| transfected construct (*mouse*) | ACOT12 shRNA-#7 | This paper |  | Lentiviral construct to transfect and express the shRNA; Targeting sequence: GCGAGGACGATCAGATATATT |
| transfected construct (*human*) | ACOT8 shRNA-#1 | This paper |  | Lentiviral construct to transfect and express the shRNA; Targeting sequence: GAGGATCTCTTCAGAGGAAGG |
| transfected construct (*human*) | ACOT8 shRNA-#2 | This paper |  | Lentiviral construct to transfect and express the shRNA; Targeting sequence: GCAGCCAAGTCTGTGAGTGAA |
| transfected construct (*mouse*) | ACOT8 shRNA-#1 | This paper |  | Lentiviral construct to transfect and express the shRNA; Targeting sequence: GGGACCCTAACCTTCACAAGA |
| transfected construct (*mouse*) | ACOT8 shRNA-#2 | This paper |  | Lentiviral construct to transfect and express the shRNA; Targeting sequence: GCTGTGTGGCTGCTTATATCT |
| antibody | anti-Flag (mouse monoclonal) | Sigma | Cat#F1804 RRID:AB_262044 | IF (1:200) WB (1:2000) |
| antibody | anti-ACOT12 (rabbit polyclonal) | Abbkine | Cat#ABP53776 | WB (1:500) |
| antibody | anti-ACOT8 (rabbit polyclonal) | Abbkine | Cat#ABP50586 | WB (1:500) |
| antibody | anti-HMGCS2 (rabbit polyclonal) | ABclonal | Cat#A14244 RRID:[AB_2761104](http://antibodyregistry.org/AB_2761104) | WB (1:1000) |
| antibody | anti-ABCD1 (rabbit polyclonal) | Abbkine | Cat#ABP54187 | WB (1:1000) |
| antibody | anti-β-actin (mouse monoclonal) | Proteintech | Cat#60008-1-Ig RRID:AB_2289225 | WB (1:2000) |
| antibody | anti-ACLY (rabbit polyclonal) | Proteintech | Cat#15421-1-AP RRID:AB_2223741 | WB (1:500) |
| antibody | anti-CPT1A (rabbit polyclonal) | Proteintech | Cat#15184-1-AP RRID:AB_2084676 | WB (1:500) |
| antibody | anti-catalase (mouse monoclonal) | Proteintech | Cat#66765-1-Ig RRID:AB_2882111 | IF (1:100) |
| antibody | anti-TOMM40 (mouse monoclonal) | Proteintech | Cat#66658-1-Ig RRID:AB_2882015 | WB (1:2000) |
| antibody | anti-LAMP2 (mouse monoclonal) | Proteintech | Cat#66301-1-Ig RRID:AB_2881684 | WB (1:2000) |
| antibody | anti-GAPDH (rabbit monoclonal) | Proteintech | Cat#60004-1-Ig RRID:AB_2107436 | IF (1:100)  WB (1:2000) |
| antibody | Acetylated-Lysine antibody (rabbit polyclonal) | Cell Signaling Technology | Cat#9441S RRID:AB_331805 | WB (1:1000) |
| antibody | HRP-conjugated goat anti-mouse IgG antibody | Thermo Fisher | Cat#A16072SAM PLE | WB (1:5000) |
| antibody | HRP-conjugated goat anti-rabbit IgG antibody | Thermo Fisher | Cat#A16104SAM PLE | WB (1:5000) |
| commercial assay or kit | Peroxisome Isolation kit | Sigma | PEROX1-1KT |  |
| commercial assay or kit | PCR-based Mycoplasma Detection Kit | Sigma | MP0035-1KT |  |
| chemical compound, drug | streptozotocin | Sangon Biotech | Cat#A610130-0100 |  |
| chemical compound, drug | ampicillin | Sangon Biotech | Cat#A610028-0025 |  |
| chemical compound, drug | streptomycin | Sangon Biotech | Cat#A610494-0250 |  |
| chemical compound, drug | tetradecanoic acid | Sangon Biotech | Cat#A600931-0250 |  |
| chemical compound, drug | sodium stearate | Sangon Biotech | Cat#A600888-0100 |  |
| chemical compound, drug | colistin | Yuanye Bio-Technology | Cat#1264-72-8 |  |
| chemical compound, drug | deuterated water (D2O) | Qingdao Tenglong Weibo Technology | Cat#DFSA180309  G100 |  |
| chemical compound, drug | sodium 3-(trimethylsilyl) propionate-2,2,3,3-d4 (TSP) | Qingdao Tenglong Weibo Technology | Cat#DLM-48-5 |  |
| chemical compound, drug | etomoxir | MedChemExpress (MCE) | Cat#828934-41-4 |  |
| chemical compound, drug | sodium palmitate | Sigma | Cat#P9767-10G |  |
| chemical compound, drug | sodium acetate | Sigma | Cat#791741-100G |  |
| chemical compound, drug | sodium 3-hydroxybutyrate | Sigma | Cat#54965-10G-F |  |
| chemical compound, drug | U- ^13^C -palmitate | Cambridge IsotopeLaboratories | Cat#CLM-6059-1 |  |
| chemical compound, drug | U- ^13^C - glucose | Cambridge IsotopeLaboratories | Cat#CLM-1396-1 |  |
| chemical compound, drug | U- ^13^C - glutamine | Cambridge IsotopeLaboratories | Cat#CLM-1822-H- 0.1 |  |
| chemical compound, drug | U- ^13^C - acetate | Cambridge IsotopeLaboratories | Cat#CLM-440-1 |  |
| chemical compound, drug | 2-^13^C- acetate | Cambridge IsotopeLaboratories | Cat#CLM-381-5 |  |
| chemical compound, drug | DAPI stain | Sigma | D9542 |  |
| software, algorithm | R | R-studio |  | R (version 3.6.3) |
