## Supplementary material for "Hepatic conversion of acetyl-CoA to acetate plays crucial roles in energy stresses": MDAR_checklist

eLife

Westbrook Centre
Milton Road
Cambridge CB4 1YG, UK

P 01223 855340
W elifesciences.org
T @elife

### Materials Design Analysis Reporting (MDAR) Checklist for Authors

The MDAR framework establishes a minimum set of requirements in transparent reporting mainly applicable to studies in the life sciences.

*eLife* asks authors to **provide detailed information within their article** to facilitate the interpretation and replication of their work. Authors can also upload supporting materials to comply with relevant reporting guidelines for health-related research (see EQUATOR Network), life science research (see the BioSharing Information Resource), or animal research (see the ARRIVE Guidelines and the STRANGE Framework; for details, see *eLife*'s Journal Policies). Where applicable, authors should refer to any relevant reporting standards materials in this form.

For all that apply, please note **where in the article** the information is provided. Please note that we also collect information about data availability and ethics in the submission form.

#### Materials:

| Newly created materials | Indicate where provided: section/figure legend | N/A |
|---|---|---|
| The manuscript includes a dedicated "materials availability statement" providing transparent disclosure about availability of newly created materials including details on how materials can be accessed and describing any restrictions on access. | Materials and methods | |

| Antibodies | Indicate where provided: section/figure legend | N/A |
|---|---|---|
| For commercial reagents, provide supplier name, catalogue number and RRID, if available. | Materials and methods | |

| DNA and RNA sequences | Indicate where provided: section/figure legend | N/A |
|---|---|---|
| Short novel DNA or RNA including primers, probes: Sequences should be included or deposited in a public repository. | Materials and methods | |

| Cell materials | Indicate where provided: section/figure legend | N/A |
|---|---|---|
| Cell lines: Provide species information, strain. Provide accession number in repository OR supplier name, catalog number, clone number, OR RRID. | Materials and methods, Generation of cell lines expressing Flag-tagged ACOT1, ACOT2, ACOT4, ACOT8, ACOT9, ACOT11 or ACOT12. Generation of cell lines for the knock down of ACOT8 or ACOT12. | |

| | |
|---|---|
| Primary cultures: Provide species, strain, sex of origin, genetic modification status. | Materials and methods,<br>Mouse primary hepatocytes were obtained from C57BL/6 mice (random sex), ACOT12 or ACOT8-knock down primary hepatocytes were induced by Adenovirus. Mouse embryonic fibroblast cell was freshly isolated from embryos of mice at 13.5 days post-coitum and further immortalized by infection of the SV-40 larger T antigen expressing retroviruses. |

| **Experimental animals** | **Indicate where provided: section/figure legend** | **N/A** |
|---|---|---|
| Laboratory animals or Model organisms: Provide species, strain, sex, age, genetic modification status. Provide accession number in repository OR supplier name, catalog number, clone number, OR RRID. | Materials and methods,<br>BALB/c and C57BL/6 mice (6-7 weeks, random sex, in groups) were obtained from Xiamen University Laboratory Animal Center (China). C57BLKS/J-$LepR^{db}$/$LepR^{db}$ (db/db) mice were purchased from GemPharmatech Co, Ltd (China). For Cre-Loxp-mediated liver-specific ACOT12 or ACOT8 knockout mice (*Acot12*$^{-/-}$ or *Acot8*$^{-/-}$), C57BL/6JGpt-*Acot12*$^{em1Cflox}$/Gpt (T039802) and C57BL/6JGpt-*Acot8*$^{em1Cflox}$/Gpt (T032253) mice were purchased from GemPharmatech Co, Ltd (China), and Acot12 $^{Flox/Flox}$ or Acot8 $^{Flox/Flox}$ mice cross with *Alb-Cre* mice (C57BL/6), confirm efficient deletion of ACOT12 or ACOT8 specifically in liver at protein (6-7 weeks, random sex, in groups). For adenovirus-mediated liver-specific RNAi mouse models, injection of 200 μL adenovirus (titer: $10^{12}$) via tail vein was performed in C57BL/6 mice (6-7 weeks, random sex, in groups) and knockdown efficiency in liver was determined by Western Blot after 2 months of injection. | |
| Animal observed in or captured from the field: Provide species, sex, and age where possible. | | N/A |

| **Plants and microbes** | **Indicate where provided: section/figure legend** | **N/A** |
|---|---|---|
| Plants: provide species and strain, ecotype and cultivar where relevant, unique accession number if available, and source (including location for collected wild specimens). | | N/A |
| Microbes: provide species and strain, unique accession number if available, and source. | | N/A |

| **Human research participants** | **Indicate where provided: section/figure legend) or state if these demographics were not collected** | **N/A** |
|---|---|---|
| If collected and within the bounds of privacy constraints report on age, sex, gender and ethnicity for all study participants. | Materials and methods,<br>Figure 1—source data 1 | |

#### Design:

| **Study protocol** | **Indicate where provided: section/figure legend** | **N/A** |
|---|---|---|
| If the study protocol has been pre-registered, provide DOI. For clinical trials, provide the trial registration number OR cite DOI. | Materials and methods | |

| **Laboratory protocol** | **Indicate where provided: section/figure legend** | **N/A** |
|---|---|---|
| Provide DOI OR other citation details if detailed step-by-step protocols are available. | Materials and methods | |

| **Experimental study design (statistics details) *** | | |
|---|---|---|
| **For in vivo studies: State whether and how the following have been done** | **Indicate where provided: section/figure legend. If it could have been done, but was not, write “not done”** | **N/A** |
| Sample size determination | Materials and methods,<br>Figure legend | |
| Randomisation | Materials and methods,<br>Figure legend | |
| Blinding | Materials and methods, | |

| | |
|---|---|
| | Figure legend |
| Inclusion/exclusion criteria | As Reviewer #3 suggested two patients with diabetes have concurrent cancer. Cancer cells have altered metabolism compared to native cells. Thus, it is possible that circulating acetate cells may be altered in these cancer patients, regardless of the presence of diabetes. And these two subjects have been taken out. |

| **Sample definition and in-laboratory replication** | **Indicate where provided: section/figure legend** | **N/A** |
|---|---|---|
| State number of times the experiment was replicated in the laboratory. | Materials and methods,<br>Figure legend,<br>More than three independent experiments | |
| Define whether data describe technical or biological replicates. | Materials and methods,<br>Figure legend,<br>More than three independent experiments | |

| **Ethics** | **Indicate where provided: section/submission form** | **N/A** |
|---|---|---|
| Studies involving human participants: State details of authority granting ethics approval (IRB or equivalent committee(s), provide reference number for approval. | Materials and methods,<br>This study was approved by the clinical research ethics committee of the First Affiliated Hospital of Xiamen University (Xiamen, Fujian, China). Before clinical serum samples acquisition, written informed consent was obtained from each patient according to the policies of the committee. | |
| Studies involving experimental animals: State details of authority granting ethics approval (IRB or equivalent committee(s), provide reference number for approval. | Materials and methods,<br>All animal studies were approved by the Animal Ethics Committee of Xiamen University (China) (acceptance no: XMULAC20190166). All surgery was performed under sodium pentobarbital anesthesia, and every effort was made to minimize suffering. | |

| | | |
|---|---|---|
| Studies involving specimen and field samples: State if relevant permits obtained, provide details of authority approving study; if none were required, explain why. | | N/A |

| **Dual Use Research of Concern (DURC)** | **Indicate where provided: section/submission form** | **N/A** |
|---|---|---|
| If study is subject to dual use research of concern regulations, state the authority granting approval and reference number for the regulatory approval. | | N/A |

#### Analysis:

| **Attrition** | **Indicate where provided: section/figure legend** | **N/A** |
|---|---|---|
| Describe whether exclusion criteria were pre-established. Report if sample or data points were omitted from analysis. If yes, report if this was due to attrition or intentional exclusion and provide justification. | | N/A |

| **Statistics** | **Indicate where provided: section/figure legend** | **N/A** |
|---|---|---|
| Describe statistical tests used and justify choice of tests. | Materials and methods,<br>Figure legend,<br>Values are expressed as mean±SD and analyzed statistically by two-tailed unpaired Student's *t* test or one-way ANOVA. | |

| **Data availability** | **Indicate where provided: section/submission form** | **N/A** |
|---|---|---|
| For newly created and reused datasets, the manuscript includes a data availability statement that provides details for access (or notes restrictions on access). | | N/A |
| When newly created datasets are publicly available, provide accession number in repository OR DOI and licensing details where available. | | N/A |
| If reused data is publicly available provide accession number in repository OR DOI, OR URL, OR citation. | NCBI Gene Expression Omnibus (www.ncbi.nlm.nih.gov/geo/):<br>GSE72086<br>GSE24207 | |

| Code availability | Indicate where provided: section/figure legend | N/A |
| --- | --- | --- |
| For any computer code/software/mathematical algorithms essential for replicating the main findings of the study, whether newly generated or re-used, the manuscript includes a data availability statement that provides details for access or notes restrictions. | | N/A |
| Where newly generated code is publicly available, provide accession number in repository, OR DOI OR URL and licensing details where available. State any restrictions on code availability or accessibility. | | N/A |
| If reused code is publicly available provide accession number in repository OR DOI OR URL, OR citation. | | N/A |

#### Reporting:

The MDAR framework recommends adoption of discipline-specific guidelines, established and endorsed through community initiatives.

| Adherence to community standards | Indicate where provided: section/figure legend | N/A |
| --- | --- | --- |
| State if relevant guidelines (e.g., ICMJE, MIBBI, ARRIVE, STRANGE) have been followed, and whether a checklist (e.g., CONSORT, PRISMA, ARRIVE) is provided with the manuscript. | | N/A |

---

* We provide the following guidance regarding transparent reporting and statistics; we also refer authors to Ten common statistical mistakes to watch out for when writing or reviewing a manuscript.

**Sample-size estimation**

- You should state whether an appropriate sample size was computed when the study was being designed
- You should state the statistical method of sample size computation and any required assumptions
- If no explicit power analysis was used, you should describe how you decided what sample (replicate) size (number) to use

**Replicates**

- You should report how often each experiment was performed
- You should include a definition of biological versus technical replication
- The data obtained should be provided and sufficient information should be provided to indicate the number of independent biological and/or technical replicates
- If you encountered any outliers, you should describe how these were handled
- Criteria for exclusion/inclusion of data should be clearly stated
- High-throughput sequence data should be uploaded before submission, with a private link for

reviewers provided (these are available from both GEO and ArrayExpress)

**Statistical reporting**

- Statistical analysis methods should be described and justified
- Raw data should be presented in figures whenever informative to do so (typically when N per group is less than 10)
- For each experiment, you should identify the statistical tests used, exact values of N, definitions of center, methods of multiple test correction, and dispersion and precision measures (e.g., mean, median, SD, SEM, confidence intervals; and, for the major substantive results, a measure of effect size (e.g., Pearson's r, Cohen's d)
- Report exact p-values wherever possible alongside the summary statistics and 95% confidence intervals. These should be reported for all key questions and not only when the p-value is less than 0.05.

**Group allocation**

- Indicate how samples were allocated into experimental groups (in the case of clinical studies, please specify allocation to treatment method); if randomization was used, please also state if restricted randomization was applied
- Indicate if masking was used during group allocation, data collection and/or data analysis
